# Blood-brain barrier dysfunction predicts cognitive trajectory after ischemic stroke

**DOI:** 10.64898/2026.02.27.708013

**Authors:** Lei Xue, Olivia A Jones, Lauren Drag, Kristy A. Zera, Li Zhu, Michael Mlynash, Natasha S. Carmichael, Chi-Hung Shu, Zuzanna Biesiada, David Seong, Owen M Thomas, Epiphani C. Simmons, Emily Huang, Kacey Berry, Philip Chung, Alperen Aslan, Rui Xu, Jarod E. Rutledge, Hamilton S.-H. Oh, Liu K. Yang, Tomin James, Marc Ghanem, Patricia Moran-Losada, Kathleen L. Poston, Victor W. Henderson, John Grainger, Stuart M Allan, Maarten Lansberg, Elizabeth C. Mormino, Tony Wyss-Coray, Alifiya Kapasi, Julie Schneider, Craig J. Smith, Laura M Parkes, Nima Aghaeepour, Marion S. Buckwalter

## Abstract

Ischemic stroke doubles the risk of dementia.^1–4^ Stroke severity and location affect cognition early,^5,6^ but late dementia risk is not related to infarct characteristics, nor is it reduced by preventing additional strokes,^3,6,7^ and its mechanism is unknown. We identified a plasma proteomic signature of chronic stroke that was consistent with blood-brain barrier (BBB) dysfunction, including a 58% decrease in plasma levels of platelet-derived growth factor B and downregulation of its pathway compared to healthy controls. During 2 years of follow-up, the stroke-specific proteome was accentuated in stroke survivors who subsequently declined in the processing speed/executive function cognitive domain. To test BBB function, we performed dynamic contrast-enhanced MRI 6-9 months after stroke in an additional cohort and found 1.7-fold higher whole brain BBB leakage compared to controls. Finally, we compared autopsy tissue from people with infarcts and dementia at death to those with infarcts and no dementia. Those who died with dementia had dramatic loss of vascular mural cell coverage compared to those without dementia (median 0.7% vs. 27%). Thus, our proteomic, functional, and structural data implicate chronic BBB dysfunction in cognitive decline late after stroke, revealing potential proteomic and imaging biomarkers and, importantly, a novel target for intervention.

## Main

Post-stroke dementia is a significant public health concern. Tens of millions of people alive today have had a symptomatic brain infarct, or stroke, and stroke at least doubles the risk that a person will subsequently develop dementia.^1,4,8–10^ Initial risk of post-stroke cognitive impairment is associated with stroke severity and location,^5,6^ but a substantial later dementia risk, which we term “infarct-induced neurodegeneration,” persists for at least a decade irrespective of patient age.^1,4^ Infarct-induced neurodegeneration is unrelated to stroke location and is not mitigated by preventing recurrent stroke (reviewed in ^2,3^), suggesting that the high risk of late cognitive decline after stroke is driven by biological processes beyond focal injury or recurrent ischemic events.

Chronic inflammation in the infarct may be one mediator of infarct-induced neurodegeneration. In a mouse model of cortical stroke, both memory and hippocampal long-term potentiation (LTP) are preserved immediately after stroke; however, mice develop cognitive deficits, progressive impairment of LTP, and aberrant neurogenesis over the subsequent months in association with lymphocytic infiltrates in the infarct.^11,12^ This progressive neurologic dysfunction is prevented by limiting lymphocyte influx into the brain.^11,13^ In small human studies, systemic immune responses to brain antigens are associated with cognitive decline,^14^ and a subset of people who have both brain infarcts and dementia have higher numbers of B lymphocytes near the infarct.^11^ To probe this and other mechanisms in the blood in an unbiased fashion, we designed the StrokeCog study^15^ to examine late cognitive decline after stroke using plasma proteomics.

### Cognitive trajectory in chronic stroke

We obtained blood, medical history, demographics, finger tapping speed, and a comprehensive cognitive battery on 124 stroke participants, all at least 5 months after ischemic stroke (**Figure 1A**). Eighty-six returned for annual cognitive batteries, with a median of 25.7 months [IQR 13.1, 35.8] between baseline and final cognitive test battery. **Extended Data Figure 1** details attrition of the study population. Participants who returned for repeat cognitive testing were similar to those who did not (**Table 1**). Tests performed and their associated domains are shown in **Figure 1B**. For each test score, raw scores were transformed to age-corrected z-scores using published norms. Z-scores for tests within each cognitive domain were averaged to create a domain-specific z-score.

**Figure 1.**
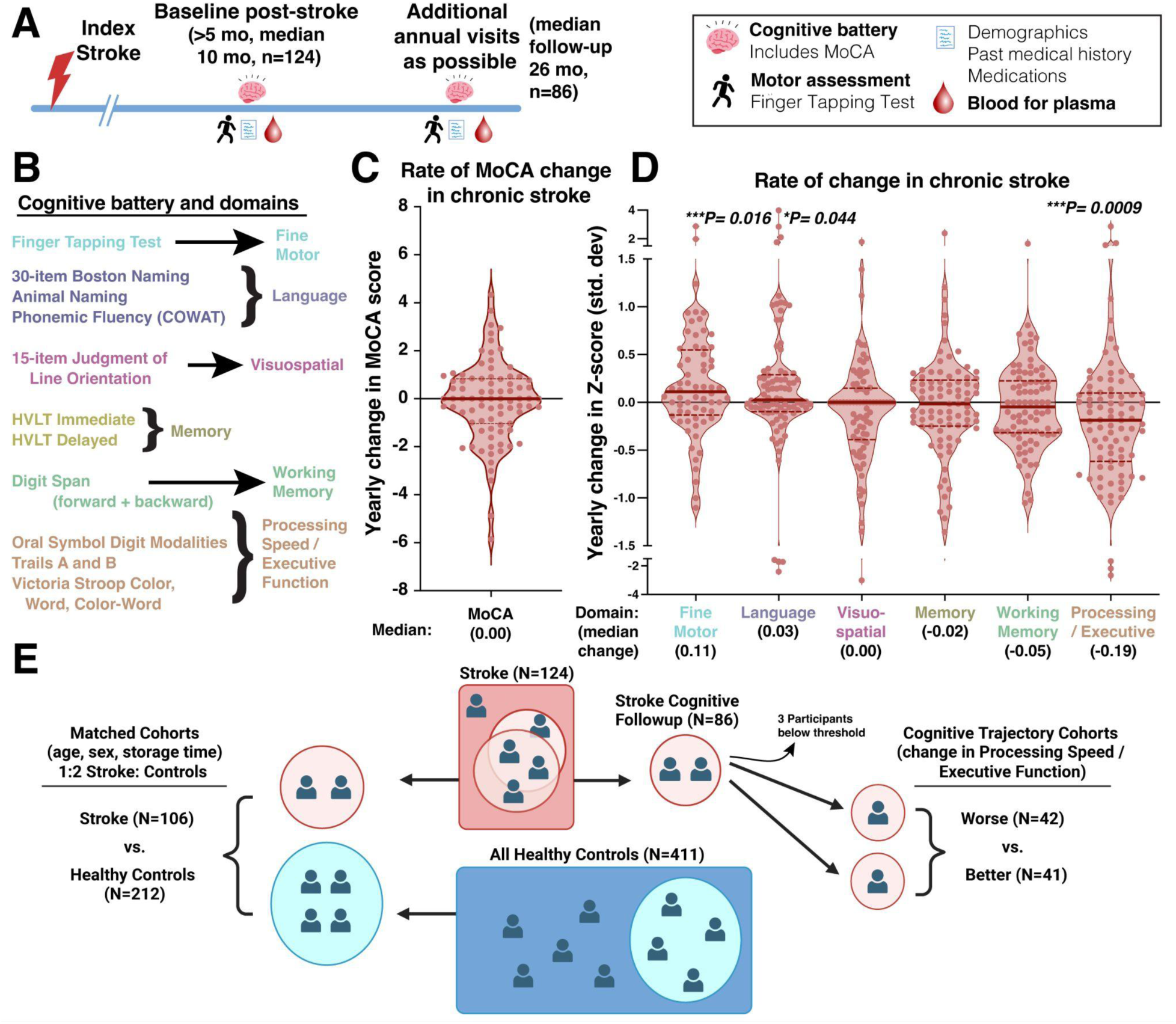
Clinical outcomes and proteomics cohort. **A)** Study design and timeline. **B)** 60-minute cognitive battery, including one motor task with available z-scores. Z-scores on individual tests were averaged to come up with Domain z-scores as indicated. Eighty-six participants returned for cognitive testing for a median of 26 months. **C)** Change in MoCA score per year. **D)** Annual change in z-scores for each domain indicates significant improvement in Fine Motor and Language with significant decline in Processing Speed and Executive Function. Wilcoxon Signed Rank test. **E)** Derivation of Proteomics cohorts. For Matched Cohorts we selected 2 controls per stroke participant, matched for age, sex, and sample storage time. To compare cognitive trajectory, we dichotomized based on the change in Processing Speed/Executive Function. Three of the 86 participants in the follow up cohort were not included because they had z-scores of −3 or lower during all of their follow-up visits, preventing accurate assessment of cognitive change.

**Table 1.** Demographic and Clinical Characteristics. “All stroke” indicates all who had plasma proteomics, “Stroke Cog. Follow-up” represents participants who returned for annual cognitive testing. Stroke Cog “better” and “worse” indicates those in the follow-up cohort whose processing speed / executive function change was better or worse than the median annual change, as indicated in Figure 1. Matched stroke and controls in the last two columns were used for the proteomic comparisons between stroke and control participants. The “All Stroke” group has 3 missing values for Education, 7 missing values for Stroke Size and 1 missing value for NIHSS. “Stroke Cog. Follow-up” group has 1 missing value for Education and 1 missing value for Stroke Size. “Stroke Cog. Better” group has 1 missing value for Education and 1 missing value for Stroke Size. “All Controls” cohort has 2 missing values for Race and 7 missing values for Education. “Matched Stroke” has 3 missing values for Education and 7 missing values for Stroke Size. “Matched Controls” group has 1 missing value for Race and 5 missing values for Education. Stroke Severity was measured by National Institutes of Health Stroke Scale, NIHSS. For Age, Education, Sample Storage Time, Stroke Size, and NIHSS, *p*-values were calculated using the Mann-Whitney U test. For Sex and Race, *p*-values were calculated using the Chi-squared test.

| Demographics | All Stroke<br>(N = 124) | Stroke Cog.<br>Follow-up<br>(N = 86) | Stroke Cog.<br>Better<br>(N = 41) | Stroke Cog.<br>Worse<br>(N = 42) | All Controls<br>(N = 411) | Matched Stroke<br>(N = 106) | Matched Controls<br>(N = 212) |
| --- | --- | --- | --- | --- | --- | --- | --- |
| Age (years) | 65 [54.8, 73.3] | 66 [55.5, 71.8]<br>$p = 0.79$ | 60 [51, 69] | 68 [60.5, 78.5]<br>$*p = 0.01$ | 70 [65, 75]<br>$p = 6.63e-7$ | 67 [58.3, 74] | 67 [63, 72]<br>$p = 0.58$ |
| Sex (% Male) | 54.84% | 58.14%<br>$p = 0.74$ | 53.66% | 61.9%<br>$p = 0.30$ | 40.39%<br>$p = 0.06$ | 51.89% | 49.53%<br>$p = 0.85$ |
| Race |  |  |  |  |  |  |  |
| White | 65.32% | 65.12% | 53.66% | 76.19% | 88.51% | 66.98% | 87.08% |
| Asian | 15.32% | 16.28% | 19.51% | 14.29% | 7.09% | 14.15% | 6.7% |
| Black | 1.61% | 1.16% | 2.44% | N/A | 2.2% | 1.89% | 2.87% |
| Hw./Pac. Islander | 0.81% | 1.16% | N/A | 2.38% | 0.24% | 0.94% | 0.48% |
| Nat. Am./ Alask. | 0.81% | 1.16% | N/A | N/A | N/A | 0.94% | N/A |
| >1 Race | N/A | N/A | N/A | N/A | 1.22% | N/A | 1.91% |
| Unknown | 3.23% | 3.49% | 4.88% | 2.38% | 0.73% | 3.77% | 0.96% |
| Other | 12.9% | 11.63%<br>$p = 1.00$ | 19.51% | 4.76%<br>$p = 0.002$ | N/A<br>$p = 0.10$ | 11.32% | N/A<br>$p = 0.15$ |
| Education (years) | 16 [14, 18] | 16 [16, 18]<br>$p = 0.22$ | 16 [16, 18] | 17 [16, 18]<br>$p = 0.27$ | 16 [16, 18]<br>$p = 4.58e-3$ | 16 [14, 18] | 16 [16, 18]<br>$p = 0.01$ |
| Sample Storage Time (days) | 969 [744, 1146] | 934 [736, 1076]<br>$p = 0.32$ | 1,035 [838, 1,119] | 806.5 [576, 988]<br>$***p = 0.002$ | 1326 [905, 1,645]<br>$p = 5.95e-14$ | 972 [792, 1,138] | 956 [679, 1,215]<br>$p = 0.80$ |
| Stroke Size (mm<sup>3</sup>) | 6 [0.9, 26.2] | 5.6 [0.9, 32.4]<br>$p = 0.99$ | 4.85 [1.1, 38.4] | 3.6 [0.69, 15.0]<br>$p = 0.15$ | N/A | 4.80 [0.9, 25.0] | N/A |
| Stroke Severity (NIHSS) | 2 [0, 6] | 2 [0, 6]<br>$p = 0.54$ | 3 [0, 7] | 2 [0, 4.75]<br>$p = 0.17$ | N/A | 2 [0, 6] | N/A |
| Compared to: |  | All Stroke |  | Stroke Cog.<br>Better | All Stroke |  | Matched Stroke |

Examining change per year, we observed overall improvement over time in finger tapping and language and no change in MoCA, visuospatial function, memory, and working memory, but a decline in processing speed/executive function (**Figure 1C&D**). Processing speed/executive function exhibited an annual z-score decline of −0.18 [IQR −0.62, −0.098]. The rate of decline was variable, and dichotomizing according to the median trajectory yielded a group with little change after baseline assessment (0.09, IQR [-0.01, 0.33], “better”) and one with substantial decline (−0.60 [-0.84, −0.36], “worse”). Risk factors for worse cognitive trajectory included older age and less diverse ethnicity but not stroke size or severity (**Table 1**). This is despite the fact that z-scores are already adjusted for age.

### Plasma proteomics in chronic stroke

We utilized aptamer-based proteomics (SomaLogic Inc., 7288 aptamers representing 6344 unique human proteins) on platelet-depleted plasma from the baseline visit in all 124 stroke participants and in 411 adjudicated healthy controls (**Figure 1E**). We selected matched stroke and healthy control participant groups for proteomic analysis based on age, sex, race, and sample storage time, resulting in 106 “matched stroke” and 212 “matched controls” (**Table 1**). Strokes were moderate to small with a median infarct volume of 4.80 mm^3^ [IQR 0.9, 25.0] and median NIHSS of 2 [IQR 0, 6]. Using cross-validated Elastic Net logistic regression, proteomics robustly distinguished stroke survivors from healthy controls with an area under a receiver operating characteristics curve (AUROC) of 0.993 (**Figure 2A**).

**Figure 2.**
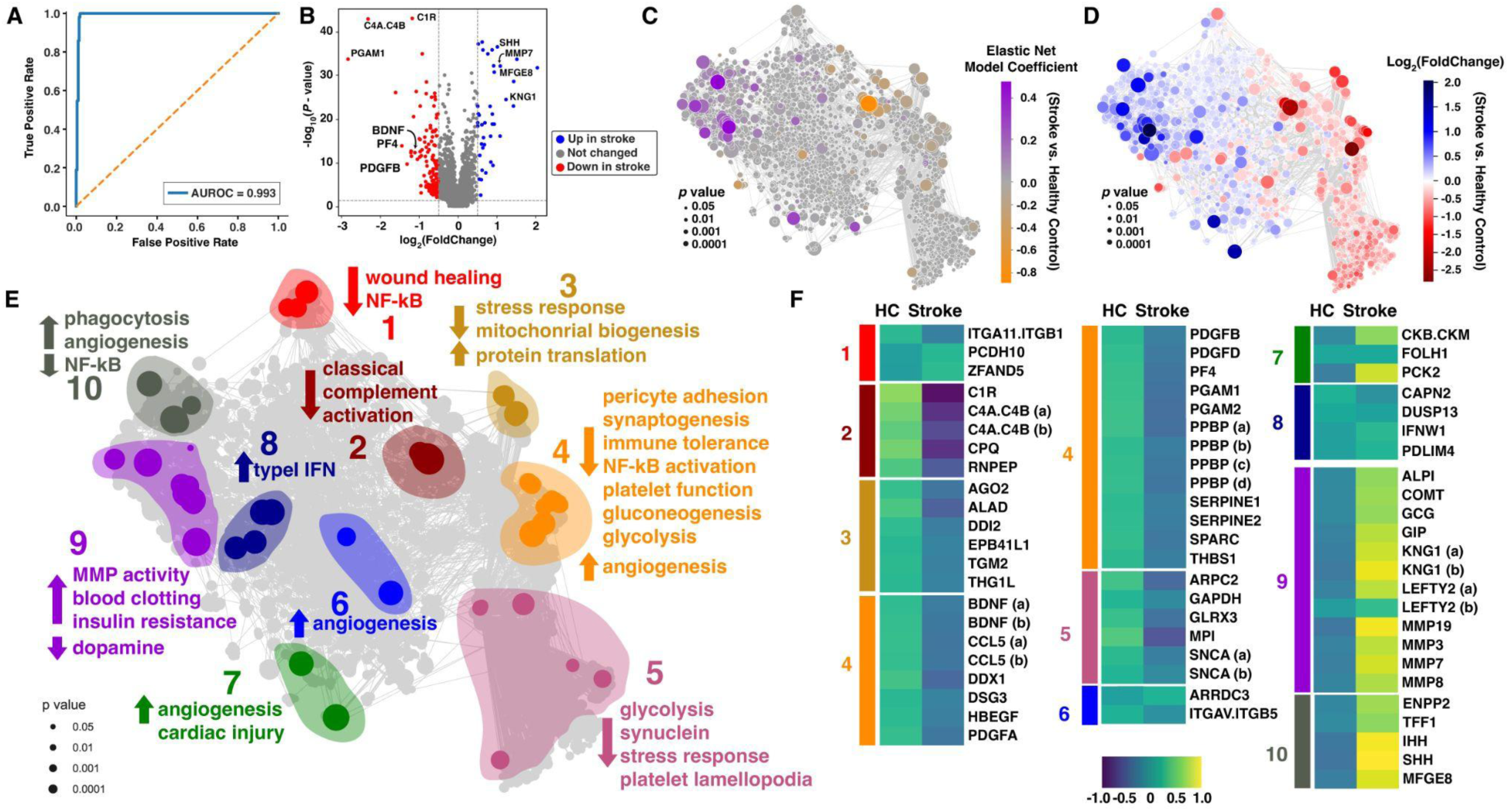
Plasma proteomics robustly discriminates chronic stroke from healthy controls. **A**) Receiver operating characteristic (ROC) curve from a participant level cross-validated Elastic Net classifier trained on plasma proteomics to distinguish healthy control (HC) and stroke groups. The model shows complete separability of the two groups (AUROC=0.993), consistent with the extensive proteomic differences observed after stroke. **B**) Volcano plot representing the plasma proteome (n=7288) differences between people with chronic stroke and healthy controls. Colored dots represent proteins with Benjamani-Hochberg adjusted *p* < 0.05 and log_2_(fold change) > |0.5| in people with strokes. Blue, upregulated; red, downregulated. **C-E)** A correlation network of the significant proteins with *p* < 0.05 calculated by Mann Whitney U test. Dot size indicates the −log10(*p*-value) and edges represent Pearson correlations > 0.5. **C)** Dot color represents the contribution, or feature coefficient, from the Elastic Net model. Positive coefficients indicate proteins with higher expression in the stroke group, while negative coefficients indicate proteins with lower expression in stroke. **D)** Dot color represents the log_2_(fold change) between stroke and healthy controls. Blue, upregulated in stroke; red, downregulated. **E)** The correlation network segregated and annotated into 10 protein functional communities, with direction and functionality noted for each community. **F)** Heat map of the proteins within the manually identified functional communities. Plasma protein levels were standardized using z-scores on log-normalized aptamer results. For proteins with more than one aptamer, they are designated here with letters, C4A.C4B (a). **Extended data Table 2** lists these aptamers using the SomaLogic identifiers for each pair.

A volcano plot visualizing differences between stroke and healthy controls (**Figure 2B**) revealed a propensity for proteins to be downregulated in chronic stroke plasma. Defining highly up- and down-regulated proteins as those with Benjamini-Hochberg adjusted *p*-values < 0.05 and log_2_(fold change) > |0.5|, we found 200 proteins were downregulated and 39 upregulated. Among down-regulated proteins were growth factors including PDGFB, BDNF, and PF4. Complement proteins C1R and C4A & C4B, but not uncleaved C4, were also downregulated, implying less activation of the classical complement pathway in stroke plasma. PGAM1, a key glycolytic enzyme, was also down-regulated. We developed a Shiny app (accessible at https://buckwalterlab.shinyapps.io/StrokeCog/) to facilitate comparisons of protein levels between stroke and healthy control groups, as well as between worse and better cognitive trajectory groups within the stroke cohort.

To visualize co-regulated groups of proteins using a pathway-agnostic approach, we generated a correlation network from the proteomic data of the 4,227 features (plasma protein levels) that were significantly different between the healthy control and stroke groups (**Figures 2C-E**). The top 20 proteins that contribute to the EN model are listed in **Extended Data Table 1**. Proteins down-regulated in stroke tended to cluster together as did those that were up-regulated, indicating this network could be used to infer functionality of proteins (**Figure 2D**).

We manually classified and annotated 10 communities containing spatially related proteins within the correlation network with larger and more significant differences between healthy controls and stroke (**Figure 2E, F**), inferring biologic effects based on published literature about protein functions and the direction of change. Cluster 4 (orange) contained the largest number of proteins and was characterized by large decreases in growth factors including PDGFB, PDGFA, and PDGFD. This analysis predicted that in people with chronic ischemic stroke there may be multiple changes relevant to the brain, including loss of pericyte adhesion, increased MMP activity, and increased angiogenesis, all associated with BBB dysfunction. The prominent decrease in PDGFB and increase in MMP7 were particularly consistent with this hypothesis. We also predict dysregulation of clotting, and decreases in wound healing, NF-kB signaling, classical complement activation, and stress response. Aptamer measurements are in theoretical units but if the targeted protein’s concentration is below the level of detection, it may appear identical. In this analysis, when aptamers targeting products of the same gene disagreed (**Extended Data Table 2**) sometimes aptamers were designed to quantify distinct protein products from one gene, or if aptamer pairs measured the same protein, typically the aptamer that showed a significant difference was more accurate (based on the company’s correlation and f-statistic measures) than the one that did not.

### Proteomics of cognitive trajectory

We compared the plasma proteome in those who would go on to have better vs. worse cognitive trajectory (**Figure 1E**, **Table 1**). The pattern of proteomic differences between the “better” and "worse” group (**Figure 3A**) was similar to the pattern of differences between stroke and healthy controls in the correlation framework (**Figure 2**), implying that a more exaggerated stroke proteomic signature predicted cognitive worsening. This was also apparent when comparing the log_2_FC for each protein between stroke vs. control and better vs. worse cognitive trajectory, there was significant correlation and the stroke-specific proteomic signature was exaggerated in the group who would go on to exhibit cognitive decline (**Figure 3B**). PDGFB was one of the most prominently downregulated proteins in both comparisons.

**Figure 3.**
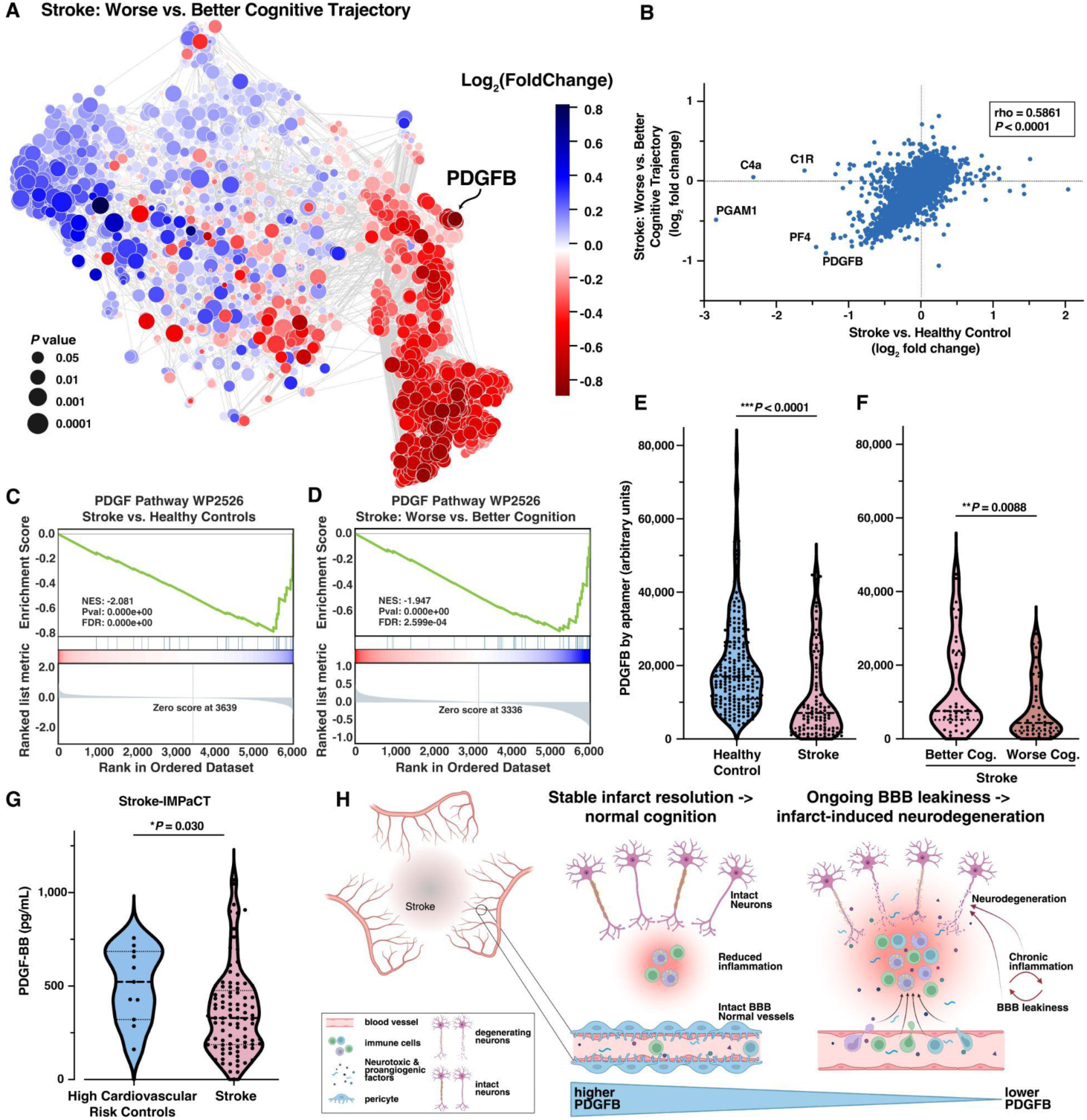
Proteomic stratification of cognitive trajectory after stroke implicates loss of BBB integrity in worsening cognition in chronic stroke. **A)** Correlation framework from Figure 2 re-drawn with data from stroke participants’ proteome, comparing worse vs. better cognitive trajectories. Dot size, −log10(*p* value); Dot color, log_2_(fold change). **B**) Scatter plot of the log_2_(fold change) of the proteins from two comparisons, worse vs. better cognitive trajectory (y-axis) and stroke vs. healthy control (x-axis). Spearman correlation. **C, D)** GSEA plot of PDGF Pathway changes in Stroke vs. healthy controls **(C),** and within stroke, worse vs. better cognitive trajectory (**D**). **E, F**) PDGFB protein levels, measured by aptamer. **E**) Matched healthy control vs. stroke participants, and (**F**) Better vs. Worse cognitive trajectory groups. **G**) PDGF-BB levels measured by ELISA in non-platelet depleted plasma from Stroke-IMPaCT participants, n = 11 high cardiovascular risk controls and 93 stroke participants 6-9 months after stroke. **H)** Hypothesis generated by the proteomic analysis, depicting how lower plasma PFGFB could promote neurodegeneration by reducing pericyte coverage on vasculature with consequent BBB dysfunction. Created by BioRender. Buckwalter, M. (2025) https://BioRender.com/f72i480

Notably, PDGFB and biologically related proteins were also prominently featured when we utilized pathway analysis tools to explore mechanistic hypotheses. Over-representation pathway analysis predicted significant changes in growth factors including PDGFB, focal adhesion and Akt pathways critical for pericyte adhesion, and immune pathways in stroke participant plasma (**Extended Data Figure 2A**), consistent with the changes we observed in our protein community analysis. Notably, about 60% of these top pathways were related to PDGFB function and BBB integrity, reflecting proteomic effects of low PDGFB protein levels in chronic stroke. Comparing stroke participants with worse vs. better cognitive trajectory, we again observed many of the same PDGFB-related pathways downregulated (**Extended Data Figure 2B**). Results were similar using Gene Set Enrichment Analysis (GSEA) (**Figure 2C, D; Extended Data Figure 3**) and consistent with decreased BBB function, immune responses, and growth factors.

PDGFB values by aptamer measurement were 58% lower in stroke than controls, and within stroke were 57% lower in those who declined cognitively after the blood draw (**Figure 3E, F**). ELISA on participant plasma confirmed that the aptamer levels correlate linearly with protein concentrations of the homodimer, PDGF-BB (**Extended Data Figure 4**). Multivariable analysis of variance conducted using a general linear model and healthy control participants with available cardiovascular risk factors (**Extended Data Table 3**) demonstrated that PDGFB levels were influenced by stroke with a strong effect, but are not affected by hypertension, diabetes, high cholesterol, sex or age (**Extended Data Table 4**). In stroke participants, the difference in PDFGB level between the worse and better trajectory groups also remained significant (*p* = 0.002) after adjustment for cardiovascular risk (hypertension, diabetes, and high cholesterol) and aspirin use. PDGF-BB was also measured in participants from the Stroke-IMPaCT study (*accepted, DOI:10.1002/alz.71261*), 6-9 months after stroke. Plasma collection in Stroke-IMPaCT was not platelet-depleted, however despite some likely contamination from platelets, PDGF-BB was 37% lower in 93 stroke participants than in 11 high cardiovascular risk controls (**Figure 3G**).

PDGFB secretion by endothelial cells is critical for both recruitment and maintenance of mural cells (largely pericytes but also vascular smooth muscle cells)^16–19^ required for proper vascular function and an impermeable BBB.^16,17,20^ Vascular mural cell adhesion requires Akt signaling,^21^ which also prominently appeared in downregulated pathways (**Extended Data Figures 2-3**). Moreover, age-related impairments occur in pericyte remodeling after injury,^22^ and this could explain why we observed more severe cognitive decline in older participants even though z-scores already account for age. We thus generated a hypothesis (**Figure 3H**) that in people with chronic brain infarction, low PDGFB secretion from endothelial cells causes mural cell loss, disrupting BBB structure and increasing BBB permeability. This BBB failure could further facilitate leukocyte invasion and leakage of plasma proteins into the damaged brain tissue, impairing neuronal function over time.

### Higher BBB leakage in chronic stroke

To test our hypothesis, we asked whether increased BBB permeability can be detected in living participants, and how commonly it occurs in chronic ischemic stroke. Fifty-four stroke survivors and 15 cardiovascular risk-matched controls from Stroke-IMPaCT (**Extended Data Figure 5** and **Extended Data Table 5**) were co-enrolled into the StrokeCog-BBB study. These participants returned for dynamic contrast-enhanced (DCE) MRI 6-9 months after their ischemic stroke to estimate the blood-to-brain volume transfer constant, *K*^trans^, a measure of BBB leakage.

Stroke survivors demonstrated spatial heterogeneity in BBB leakage, with discrete hotspots of high *K*^trans^ in and around the stroke lesion, and more rarely in the otherwise normal-appearing tissue remote from the infarct. In some participants, we observed lower *K*^trans^ resembling control brains (**Figure 4A-B)**. Median whole brain *K*^trans^ was 0.179 x10^-3^ min^-1^ [IQR 0.133, 0.210] in the stroke survivors, 1.7-fold higher than in controls, where median whole brain *K*^trans^ was 0.103 x10^-3^ min^-1^ [IQR 0.037, 0.125] (**Figure 4C**), demonstrating persistently elevated BBB leakage in chronic ischemic stroke compared to individuals with a significant cardiovascular risk burden but no stroke. The majority of stroke survivors (74%) had whole brain *K*^trans^ higher than the upper quartile of controls. Those with higher leakage had a greater prevalence of hypertension (*p* = 0.03, **Extended Data Table 5**), but no difference in demographics, vascular risk factors, baseline stroke characteristics, or cognitive performance.

**Figure 4.**
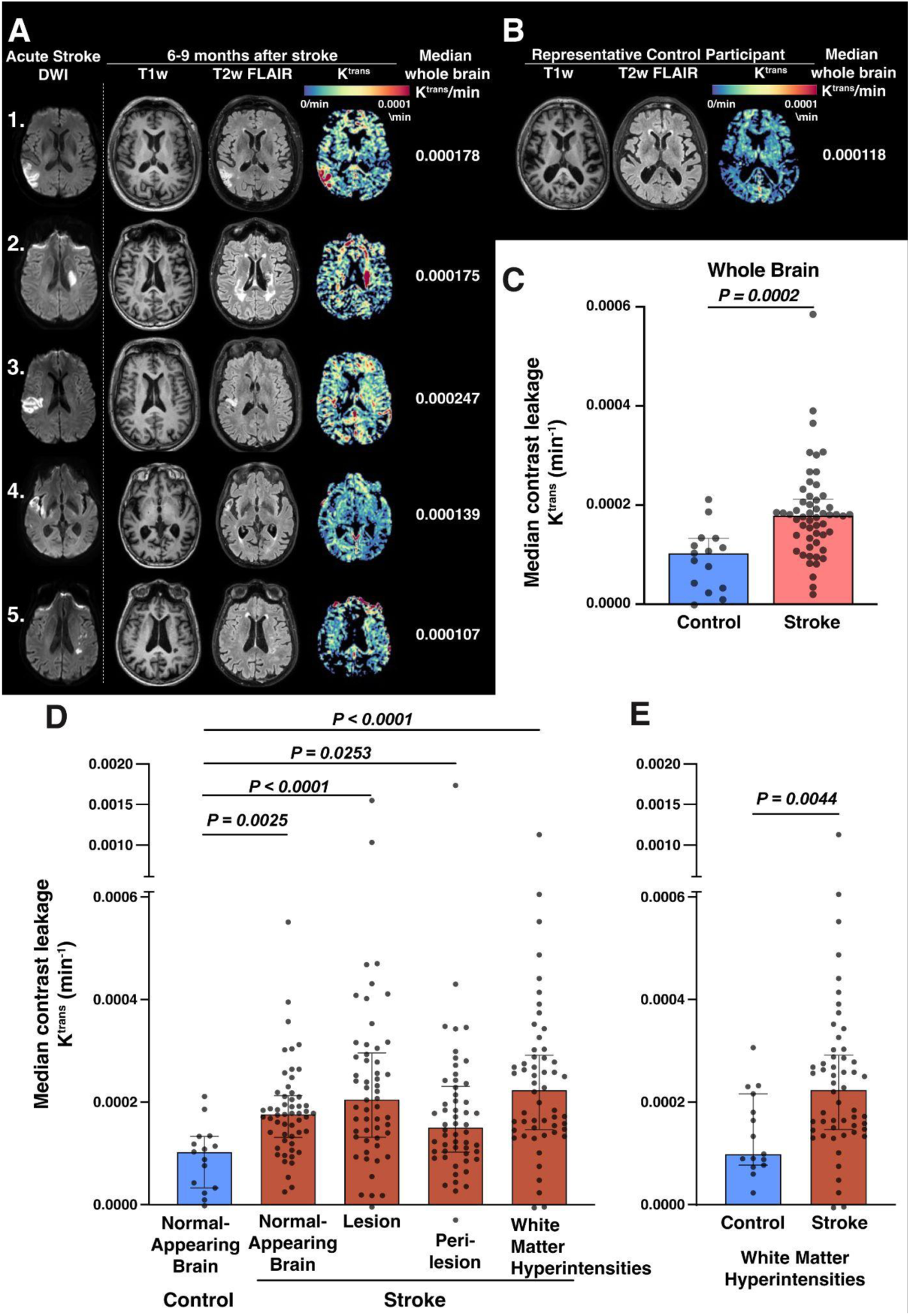
DCE-MRI of participants with chronic stroke demonstrates increased BBB leakage compared to high cardiovascular risk controls. Fifty-four stroke participants from Stroke-IMPaCT underwent MRIs during the acute stroke to assess lesion location and size, then 6-9 months later with DCE-MRI to estimate BBB leakage using median whole brain *K*^trans^. These were compared with 15 high cardiovascular risk control participants. **A)** For 5 representative stroke participants, the acute diffusion weighted image (DWI) highlighting the index infarct is presented alongside images collected at 6 – 9 months after stroke, including a structural *T*_1_- weighted image, a *T*_2_-weighted FLAIR image, and the *K*^trans^ map from DCE-MRI. Median whole brain *K*^trans^ estimates for the pictured participants are annotated to the right of the *K*^trans^ map. **B)** For a representative control participant, structural *T*_1_-weighted and *T*_2_-weighted FLAIR images are presented alongside the *K*^trans^ map from DCE-MRI. **C)** Comparison of whole-brain *K*^trans^ in the high cardiovascular risk control group versus the stroke group at 6 – 9 months after onset. *p*-value, Mann Whitney test. **D)** Segmentation of DCE-MRI scans demonstrates that all regions in stroke participants have higher median *K*^trans^ estimates than the normal-appearing tissue of controls. *p*-values, Kruskal-Wallis non-parametric ANOVA with Dunn’s post-hoc test comparing to control. **E)** Median *K*^trans^ in the white matter hyperintensities of stroke survivors is higher than in the white matter hyperintensities of non-stroke controls. *p*-value, Mann Whitney test; Bars, median and IQR.

BBB leakage was also higher within all regions (normal appearing tissue, acute stroke lesion, peri-lesion, and white matter hyperintensities) in the stroke brains than in normal appearing tissue in control brains (*p* = 0.0002, Kruskall-Wallis nonparametric ANOVA, **Figure 4D**). We saw no consistent differences across the cohort between BBB leakage in the normal-appearing tissue and that in the stroke lesion or peri-lesion within the same brain, however in some individuals there were marked increases and decreases in these regions, particularly in the stroke lesion (**Extended Data Figure 6**). Finally, in the white matter hyperintensities of stroke brains, which exclude the stroke lesion, we found 2.3-fold higher *K*^trans^ compared to the white matter hyperintensities of control brains (**Figure 4E**).

### Mural cell loss in chronic stroke

To further test our hypothesis that there is vascular mural cell loss, BBB leakiness and cognitive decline, we selected brain sections from well-characterized cohorts at the Rush Alzheimer’s Disease Center with extensive (average 8 years) pre-morbid cognitive data and detailed neuropathological characterization. We selected 8 controls with neither infarcts nor dementia, and two groups of 26 participants each with chronic infarcts that were defined pathologically. Of the two infarct groups, one had no dementia at the time of death, and the other had dementia, based on a clinical consensus of cognitive status prior to death (**Extended Data Table 6**). Among cases with pathologic infarcts without dementia, 54% had mild cognitive impairment (MCI) and 46% had normal cognition. Pathologically, these cases had minimal possible scores for AD pathologies within the larger dataset and had similar severity levels of vessel pathologies with the exception of atherosclerosis, which was greater in the group with infarcts and dementia and also in the control group. Among participants with infarcts, the dementia and non-dementia groups did not differ in the number of subcortical lacunar infarcts or in the frequency of gross or microinfarcts.

We stained for an endothelial cell marker (CD31) and a mural cell marker (PDGFRB, the receptor for PDGFB). Median vascular coverage by mural cells was 63.0% [IQR 53.3, 85.5] in controls and not different in the infarct and no dementia group at 27.5% [IQR 7.0, 70.5] (**Figure 5A-B**). However, those with dementia and infarcts exhibited significantly lower mural cell coverage at 0.7% [IQR 0.0, 12.5]. This remained true after adjusting for age at death, sex, and atherosclerosis score (*p* = 0.005, Quade nonparametric ANCOVA), consistent with our hypothesis that mural cell loss is associated with dementia in those with chronic brain infarctions.

**Figure 5.**
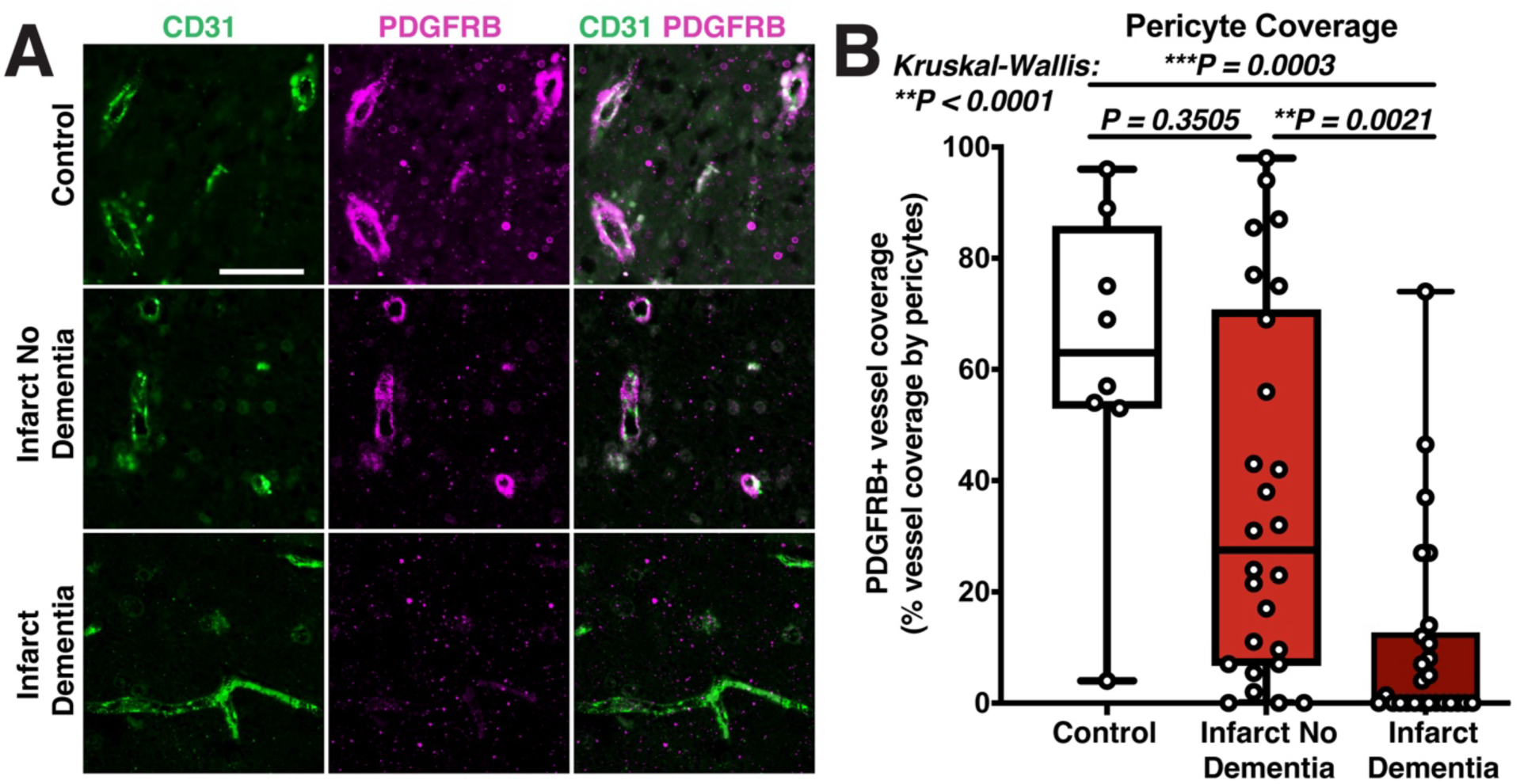
Mural cell loss in autopsy sections. **A)** Representative photomicrographs of immunofluorescent stains for CD31 (green) and PDGFRB (magenta), as well as merged images. **B**) Quantification of mural cell coverage in the 3 groups with n=8 for controls and n=26 for each infarct group. Bars, IQR with range; *p*-values, Kruskal Wallis ANOVA with Tukey post-hoc.

Finally, given the involvement of lymphocytes in the mouse model pathology^11,13^ we tested whether B or T lymphocyte numbers in brain tissue differed between groups. Neither CD20-expressing B lymphocytes or CD3-expressing T lymphocytes were different between controls and those with infarcts with or without dementia (**Extended Data Figure 7A-B**). Mural cell coverage did not correlate with the number of either type of lymphocyte (Spearman *rho* and *p*, −0.19 and 0.14 for B lymphocytes and −0.069 and 0.60 for T lymphocytes). However, there was an association between CD20 density and dementia by binary logistic regression (*p* = 0.042, OR= 1.16 (95% CI 1.01-1.35)), suggesting that lymphocytic neuroinflammation and BBB leakiness may promote cognitive decline independently.

## Discussion

We report here that a comprehensive cognitive battery detects declines in processing speed and executive function within a few years after stroke. Unbiased high-density plasma proteomics identified a chronic ischemic stroke plasma proteomic signature predicting mural cell loss and BBB breakdown. This signature both preceded and was more pronounced in participants with subsequent cognitive decline in our StrokeCog cohort. Then, we confirmed high BBB leakage on DCE-MRI in a second independent cohort of chronic stroke survivors. Finally, we demonstrated that the structural BBB changes predicted by our proteomics were present in a third independent autopsy cohort, finding specifically that there is dramatic mural cell loss adjacent to cerebral infarcts in those that die with dementia but not in cases with no premorbid dementia.

Our comprehensive cognitive battery revealed that cognitive decline begins within the first few years after stroke, with processing speed/executive function decreasing nearly 0.2 standard deviations annually across a sample that already exhibits a baseline level of cognitive impairment.

Participants in the bottom half, i.e. our “worse trajectory” group, declined even faster, at a median of 0.6 standard deviations per year over the two-year follow-up alone. Because we are examining change in z-scores that are already adjusted for age, the increased age in the “worse trajectory” group may reflect increased vulnerability to injury in aged individuals. This adds to prior studies of people with stroke that used cognitive screens or medical record searches to define dementia,^1,4,6,23^ and corroborates findings that aging and stroke impact processing speed.^23,24^

We hypothesized that BBB leakage due to mural cell loss predicts cognitive decline after completing an unbiased analysis of plasma proteomic data from 7288 proteins in a moderately large sample size of 106 stroke and 212 matched healthy control participants. An elastic net model robustly distinguished the stroke group from controls, with an accentuated stroke-specific proteomic signature in those with worse cognitive trajectory. Using both correlation network-defined protein communities and pathway level analyses, we predicted decreased numbers of pericytes and other mural cells, and increased BBB leakage in the cognitively worsening group. In addition to prominently decreased PDGFB, the signature included protein changes consistent with BBB leakage including increased MMP7, shown to correlate with BBB dysfunction in traumatic brain injury,^25^ and downregulated Akt signaling, crucial for pericyte adhesion.^21^

We then verified that BBB leakage persists in stroke survivors 6-9 months after their strokes using DCE-MRI. There was higher *K*^trans^ in stroke survivors compared to high cardiovascular risk control brains, in all regions including the chronic stroke lesion, peri-lesion, and normal-appearing tissue. *K*^trans^ was also higher in the white matter hyperintensities of stroke survivors compared to those in controls. Stroke survivors with more BBB leakage had a greater incidence of hypertension, implicating underlying hypertension as a pathologic factor in chronic BBB dysfunction after stroke. Our findings extend the timeline of prior studies that showed elevated *K*^trans^ at ∼3 months after lacunar stroke in normal appearing white matter^26,27^ compared to control participants, and in white matter hyperintensities compared to normal-appearing tissue.^28^

We next evaluated autopsy samples from a third cohort to determine whether mural cell loss occurs in individuals with both chronic cerebral infarcts and dementia diagnosis proximate to death. We utilized a longitudinal community cohort enrolled without dementia that has undergone extensive cognitive and neuropathological characterization. Mural cell loss adjacent to infarcts was profound in those with dementia but absent in those with infarcts alone, where coverage was not different from healthy controls. There were no differences in people with microscopic vs. lacunar vs. large vessel strokes (data not shown), suggesting that the same pathology likely occurs in clinically silent infarcts. This aligns with findings from our proteomics cohort and published studies showing infarct-induced neurodegeneration independent of infarct size.^6,29^ Cases from this autopsy cohort were selected to assess vascular cases with minimal AD pathology, and the younger age of our proteomics cohort compared to typical memory study participants likely reflects similarly low AD pathology.

Our analysis of autopsy samples was limited to tissue adjacent to infarcts, however vascular mural cell loss may be diffuse and present throughout the brain. We found here increased *K*^trans^ in chronic stroke across the whole brain, even after excluding the infarct and white matter disease. In our mouse model of infarct-induced neurodegeneration, stroke alone is sufficient to induce chronic loss of pericytes and BBB leakage,^13^ suggesting that mural cell loss and subsequent BBB leakage may be caused by the infarct alone in people as well. In addition, both ApoE4 and aging lead to pericyte vulnerability in mice,^22,30^ so could exacerbate BBB dysfunction in people with stroke. Similar pathology has also been associated with AD and ApoE4 in some studies.^31–33^ However, stroke could be a key underlying pathology there as well, as ApoE4 status is also a strong risk factor for stroke^34^ and approximately 60% of clinically-identified AD cases have infarcts at autopsy.^35^ Notably, stroke participants were included in studies initially reporting BBB leakage in humans with AD and ApoE4,^31,32^ so infarct-induced neurodegeneration may have contributed to those dementia diagnoses. It may also be that brain infarction, age- and ApoE4-induced vascular vulnerability, and AD pathology can all damage the BBB through similar or identical mechanisms. Integration of these data into multivariate and machine learning models^36^ will be needed to disentangle the effects of infarct-induced neurodegeneration from those of AD and other common neuropathologies.

It is clear that robust biomarkers of BBB dysfunction in stroke survivors are required. We show that PDGFB is low in chronic stroke, and even lower in the “worse” cognitive trajectory group. In addition to its key role in maintaining the BBB, PDGFB is synthesized by platelets, and thus measurements vary depending on plasma preparation. Indeed, here we observed lower PDGFB levels in platelet-depleted plasma from our initial proteomic study than in conventionally prepared plasma from the Stroke-IMPaCT study. Nonetheless, we found that PDGFB was decreased in stroke survivors compared to high cardiovascular risk controls and does not change with cardiovascular risk (hypertension, hyperlipidemia, or diabetes), and conventional preparation is more clinically translatable. Ongoing cognitive follow-up in Stroke-IMPaCT will determine whether PDGFB measured in conventionally prepared plasma predicts cognitive decline. If validated, future studies should examine PDGFB kinetics over time, and its ability to predict long-term cognition 5-10 years after measurement.

The complete plasma proteomic chronic stroke signature may be an even better biomarker of cognitive risk after stroke, particularly when paired with routinely collected clinical data. Recent advances in multimodal machine learning frameworks that integrate high-dimensional -omics data with longitudinal electronic health record information demonstrate improved patient stratification and predictive performance, even in modestly sized cohorts, providing a clear pathway for translating post-stroke proteomic signatures into clinically actionable prediction models.^37^ Finally, BBB leakage measured with DCE-MRI could serve as an important biomarker of cognitive decline if sufficiently sensitive and specific. This work is ongoing in the NIH-NINDS funded StrokeCog-BBB study (R01 NS124927), which will test whether BBB dysfunction 6-9 months after stroke predicts cognitive decline several years later. While isolated infarcts remain unexamined, other studies in small vessel disease that include stroke survivors are promising, with BBB leakage in white matter hyperintensities showing an association with cognitive decline at 2 and 1 year follow-up, respectively.^38,39^ Interestingly, we observed here that *K*^trans^ was higher in white matter hyperintensities in people with stroke compared to white matter hyperintensities of controls. This may be related to ongoing Wallerian degeneration in and around areas of infarction, and/or could be a sign that stroke amplifies autoimmune responses against the brain that then affect all regions.

Indeed, our prior work highlighted B-cell driven adaptive immunity as a mechanism of infarct-induced neurodegeneration, because ablation of B lymphocytes prevents post-stroke cognitive decline in mice, and B lymphocytes accumulate in infarcts in both mice and people.^11^ In mice, lymphocytic infiltrates and BBB integrity both improve when VLA-4 or VCAM1 is blocked.^13^ In humans, autoimmune responses do occur after stroke in association with worse outcomes, including cognition.^14,40,41^ We previously reported worse cognitive decline in people with increased activation of classical monocytes 2 days post-stroke, a time when circulating brain antigens are present.^42^ This is consistent with a model where the infarct generates brain antigens and autoimmune responses are more likely when the peripheral monocytes are stimulated to provide an adjuvant-like effect. The work here also demonstrated an association between B lymphocyte density in infarcts and cognitive trajectory. Low systemic PDGFB may reduce mural cells on new and established vasculature, leading to structural BBB deficits that drive leakage of plasma proteins and immune cells, perpetuating chronic inflammation. We did not detect this interaction here but may have been underpowered, or immune responses may wane earlier than BBB dysfunction and not be as prominent at autopsy.

## Conclusion

We utilized three complementary techniques in three independent cohorts of stroke survivors, and all confirmed chronic BBB dysfunction. The proteomic evidence of BBB dysfunction precedes worsening cognition, and the loss of mural cells at autopsy is tightly linked to cognitive outcome. A causal role of BBB dysfunction in post-stroke cognitive decline is consistent with evidence in mice where blocking endothelial activation with either an anti-VLA4 or anti-VCAM1 antibody improves mural cell coverage, decreases BBB leakage and lymphocyte infiltration, and prevents cognitive decline.^13^ Thus, among other immune therapies, FDA-approved anti-VLA4 antibody treatment, which limits BBB leakage and improves cognition in multiple sclerosis,^43–47^ may also prevent mural cell loss, BBB dysfunction, and cognitive decline in people with chronic ischemic stroke. A similar or greater magnitude of late cognitive decline is also observed after hemorrhagic stroke and subarachnoid hemorrhage, suggesting that chronic BBB dysfunction may be a common therapeutic target across multiple types of stroke.^4,29^ Given that there are 12 million strokes worldwide with 5 million survivors annually,^9^ future therapies that prevent or treat BBB leakage could mitigate stroke-induced cognitive disability in up to 2.5 million people each year.

## Methods

### Cohorts

#### Proteomics Cohort

Plasma proteomics was performed on participants recruited at Stanford as part of the Stroke-CyTOF^42^ and StrokeCog studies.^15^ We included people with ischemic stroke at least 5 months after stroke. Participants with pre-existing dementia, immune disorders requiring medication, or life expectancy of less than one year were excluded. Healthy control proteomic data were from three observational studies: the Stanford Alzheimer’s Disease Research Center (ADRC, N=194), the Stanford Aging and Memory Study (SAMS, N=192), as well as from additional affiliated studies focused on Alzheimer’s disease or Parkinson’s disease (N=25). Healthy control status was determined using study specific criteria.^48,49^ All blood was processed identically and proteomics run together at SomaLogic, Inc (Boston, MA).

#### Stroke-IMPaCT and StrokeCog-BBB

The Stroke-IMPaCT study (*accepted, DOI:10.1002/alz.71261*) was designed to parallel StrokeCog in terms of cognitive testing. Timepoints are similar except for the addition of an acute timepoint. The study was conducted in full conformity with the current revision of the Declaration of Helsinki, with approvals obtained from the Health Research Authority (IRAS ID: 275726), and written informed consent was obtained from all participants; consent was obtained from a personal consultee if a potential participant was deemed lacking capacity to provide written informed consent due to a reduced level of consciousness, or problems with cognition or communication. Participants were recruited from the Comprehensive Stroke Centre in the Manchester Centre for Clinical Neurosciences at Salford Royal Hospital (Salford, UK) during their inpatient stay following acute ischemic stroke and underwent MRI at 6-9 months after stroke. Patients with symptomatic ischemic stroke as confirmed by routine brain imaging, over the age of 45 years, able to undergo venous blood sampling within 96h of stroke symptom onset, willing to consent to study participation and follow-up, sufficiently fluent in written and spoken English, and living in the Greater Manchester area or willing to return for follow-up were included. Exclusion criteria were: a score of 2 points or more on the language component of the NIHSS; pre-existing neurological, psychiatric, or other condition (e.g. blindness) that would make it difficult to accurately assess neurologic and/or cognitive outcome; history of hemorrhagic stroke (prior to ischemic stroke) at the time of enrolment; unlikely to survive to follow-up or palliative care considered to be imminent; participation in a clinical trial of a medicinal product that might affect inflammation, or an established diagnosis of dementia predating the stroke. Routine clinical brain MR images performed as part of normal clinical care from within one month of stroke onset were obtained where available.

A group of age and risk matched controls with no history of stroke or dementia were identified through the National Institute for Health and Care Research (NIHR) Research for the Future programme, and underwent identical assessments. Inclusion criteria were an age of above 45 years and at least two of the following vascular risk factors: treated hypertension, diabetes mellitus, treated hyperlipidaemia, current smoker, confirmed atrial fibrillation, confirmed left ventricular hypertrophy, or raised body mass index >25. Exclusion criteria were history of any previous stroke, transient ischemic attack, significant head injury, autoimmune neurological disease, current brain tumour, current nervous system infection or neurodegenerative disease; treatment for autoimmune, oncological or infectious disease within the preceding six weeks; pre-existing neurological, psychiatric, or other condition that would make it difficult to accurately assess cognitive outcomes; current participation in a clinical trial of a medicinal product or device; acute ischemic event or major surgery within the preceding three months; or a score of less than 18 on telephone MoCA.

Stroke-CogBBB at The University of Manchester is a sub-study of Stroke-IMPaCT that began in May 2022. For this sub-study, Stroke-IMPaCT participants underwent DCE-MRI at 6-9 months after the enrollment stroke. Inclusion into this sub-study required that they had an MRI demonstrating acute infarct and could safely undergo MRI with gadolinium-based contrast, thus participants with contraindication to MR scanning, claustrophobia, or an estimated glomerular filtration rate <30 ml/min/1.73m^2^ in three months prior to scanning were not approached. In total, 58 returned for DCE-MRI assessment at a median of 6.9 [IQR 6.5, 7.5] months post-stroke. Four stroke participants were excluded due to incomplete scans (n=2), absent acute infarct on initial imaging (n=1), and poor data quality (n=1). Among 17 recruited controls who completed the research MRI protocol, two were excluded as a result of incidental findings (tumor, n=1; old lacunar infarcts, n=1). Thus, DCE-MRI analysis included 54 stroke survivors and 15 cardiovascular risk-matched controls.

#### Autopsy cohort

Autopsy brain samples were obtained from Rush University Alzheimer’s Disease Center Pathology Core-/Rush Memory and Aging Project (MAP) and Religious Orders Study (ROS).^50,51^ Brain samples were selected to be matched and minimized for AD pathology. Twelve 6-micron thick sections were cut from paraffin blocks containing infarcts for both infarct groups, and from cortical sections for controls. Infarcts were confirmed by a neuropathologist who examined an H&E stain of sections 1 and 12, i.e. both sides of the sections to be immunostained, to ensure that each section contained an infarct prior to shipping to Stanford.

### Cognitive testing

This was performed as described^15^ for the StrokeCog cohort, with 5 participants in the follow-up cohort recruited from the earlier prospective cohort Stroke-CyTOF.^42^ For those participants, their first cognitive battery was largely identical but had a different version of the Stroop test, and domain z-scores were calculated similarly. The cognitive battery for StrokeCog and Stroke-IMPaCT consists of 9 standardized neuropsychological tests yielding 13 cognitive variables. Variables were grouped into the following cognitive domains based on a pair-wise undirected Pearson correlation graph (t-distributed stochastic neighbor embedding [tSNE] plot): language (30-item Boston Naming Test, controlled oral word association test [COWAT], animal naming); processing speed/executive functioning (Oral Symbol Digit Modalities Test [SDMT], Trails A and Trails B, Victoria Stroop Dot, Victoria Stroop Word, Victoria Stroop Color-Word); visuospatial functioning (15-item Judgment of Line Orientation [JLO]), working memory (Wechsler Adult Intelligence Scale-III Digit Span); and memory (Hopkins Verbal Learning Test-Revised [HVLT] Immediate Recall, HVLT Delayed Recall). A 30-minute battery consisting of a subset of tests was also examined based on the NINDS/CSN harmonization standards (NINDS/CSN-30): Trails A, Trails B, SDMT, animal naming, COWAT, HVLT Immediate Recall, and HVLT Delayed Recall. For individual tests, raw scores were transformed to age-corrected (and in some instances education- and sex-corrected) z-scores based on normative data provided by the test publisher or available in other published sources. To minimize the potential impact of extreme scores on calculations, z-scores were truncated at a minimum of z = −3 and a maximum of z = 3. Composite domain scores were calculated for each cognitive domain by averaging the z-scores for individual tests within each domain

### Blood processing

Plasma samples were collected from participants at Stanford (proteomic cohorts, including healthy controls) in ethylenediaminetetraacetic acid (EDTA) tubes that were processed for platelet-depleted plasma as follows. They were immediately placed on ice prior to processing. Samples were centrifuged at 2000 × g for 10 min at 4 °C with the brake set to level 1. Plasma was carefully collected from the upper two-thirds of the supernatant to avoid disruption of the buffy coat. Plasma was gently mixed, aliquoted and stored at −80 °C until analysis.

Conventional plasma samples were obtained at Manchester for StrokeCog-BBB stroke participants and high cardiovascular risk controls. Venous blood was drawn from stroke participants within 96 h of stroke symptom onset or time last seen well. Whenever possible, blood draws were performed between 0800 and 1400. Blood was collected in EDTA tubes and processed within 4 h. Plasma was isolated by centrifugation of EDTA blood tubes at 2000 g for 10 min at 4⁰C. Plasma was aliquoted and stored at −80°C until analysis.

### Proteomic methods

Plasma protein profiling was performed using the SomaLogic SOMAScan platform (SomaLogic Inc.). This technology employs slow off-rate modified DNA aptamers (SOMAmers) to quantify the relative abundance of 7,288 human proteins within plasma with high specificity. The platform’s performance characteristics have been previously described.^52,53^

Samples were stored at −80 °C and shipped on dry ice to SomaLogic. To control for intra- and inter-batch variability, control, calibrator, and buffer samples were included in each 96-well plate. Sample data were normalized to a pooled reference using an adaptive maximum likelihood procedure. Samples with signal intensities deviating significantly from expected ranges were flagged by SomaLogic.

Data normalization was performed by the manufacturer in three stages. First, hybridization control probes were used to adjust for individual sample variance (Hybridization Control Normalization). Second, inter-sample differences within each plate were corrected through median normalization (Intraplate Median Signal Normalization). Finally, plate scaling and calibration were applied to remove variability across assay runs (Plate Scaling and Calibration)

### Proteomic Analysis

The cohort of stroke and control patients was matched in a 1:2 ratio for age, sex and plasma sample storage time using the cardinality method by the R package, *MatchIt.*^54^ Other organisms except for human proteins were removed from the SomaScan proteomics data. We developed an Elastic Net regression model on the proteomics data to predict the Health Control and Stroke groups^55^. The input matrix to the model was standardized, and the model was executed with the hyperparameter setting ‘l1_ratio=0.5’ to optimally balance the contribution of L1 and L2 regularization. Model performance was evaluated using stratified 5-fold cross-validation, with class proportions preserved in each fold. All preprocessing steps, including feature scaling, were performed within each training fold to prevent information leakage. The feature importance in the Elastic Net model was determined by the magnitude of the coefficients assigned to each protein. The *p*-value of the predicted probability between the treatment groups was calculated by *t*-test. The significantly altered proteins between Health Control and Stroke groups were identified using the Mann-Whitney U test (*p*<0.05) for Over-Representation analysis. For Gene Set Enrichment Analysis (GSEA), proteins were pre-ranked based on *p*-values from the univariate test. The pathway analysis was performed using the Gene Ontology (GO) pathway^56,57^, WikiPathways^58^ and Kyoto Encyclopedia of Genes and Genomes (KEGG) databases^59^, facilitated by the *GSEApy*^60,61^ toolkit. Correlation networks were generated by *Networkx*^62^ and laid out by the t-SNE algorithm. The key proteins of these annotated communities were normalized by standardizing each feature to have a mean of zero and a standard deviation of one in the heatmap. This standardization process ensures that each feature contributes equally to the model by removing any inherent biases due to differences in scale. All analyses were performed with Scipy, Numpy, Seaborn and Scikit-learn in Python v2.7.16^63–65^.

### ELISAs

ELISAs were performed on plasma samples to measure PDGF-BB, the homodimer of PDGFB. We used either conventional ELISA (on Stanford samples) or the Ella Automated Immunoassay System (Bio-Techne, Minneapolis, USA) (on Manchester samples). Conventional ELISA was performed in duplicate on plasma samples with the Quantikine Human PDGF-BB ELISA kit (DBB00; R&D Systems, Minneapolis, MN, USA) according to the manufacturer’s instructions. For the ELLA Automated Immunoassay System plasma was diluted in sample diluent (Cat No. 896098, Bio-Techne) according to the recommended dilution factor for that analyte, and loaded into Ella cartridges. Wash buffer (Cat No. 896055, Biotechne) was loaded into the necessary wells, and cartridges were run on the Ella platform.

### DCE-MRI Scans

DCE-MRI was performed on StrokeCog-BBB participants in Manchester, England, at 6 – 9 months after the stroke, or at enrollment for controls. Participants with no contraindication to MR scanning, no known renal impairment based on an estimated glomerular filtration rate < 30 ml/min/1.73m² in three months prior to scanning, and no known claustrophobia underwent the DCE-MRI scan.

#### Processes and model definitions

All DCE-MRI quantities, processes and model definitions are OSIPI CAPLEX compliant.^66^ CAPLEX definitions can be accessed by clicking on quantity, process or model hyperlinks.

#### MRI acquisition

MRI data were acquired on a Philips Ingenia Elition X 3T scanner (Salford Royal Hospital) with a 32-channel head coil. A dynamic series of 3D T_1_-weighted Fast Field Echo (T_1_-FFE; spoiled gradient recalled echo) images were acquired with a spiral k-space acquisition mode (spiral out, ‘stack of spirals’ 3D distribution type, 10 spiral interleaves, acquisition window of 10 ms), a <u>repetition time</u> of 10.6 ms, <u>echo time</u> of 0.9 ms, <u>prescribed excitatory flip angle</u> of 12 degrees, an in-plane resolution of 1.5 mm x 1.5 mm, 70 contiguous axial slices of 2 mm thickness, and an acquisition time of 7.6 s for each dynamic. One hundred and fifty-seven dynamic images were acquired over 20 minutes. On the 8^th^ dynamic, a gadolinium-based contrast agent (Gd-DOTA; Dotarem, Guebert, France) bolus was administered using a power injector at a dose of 0.1 mmol/kg and an injection speed of 1 ml/s.

Prior to the dynamic scan, a series of additional 3D T_1_-FFE images were acquired at 4 different <u>excitatory flip angles</u> (2, 6, 10, and 15 degrees) to calculate the <u>pre-contrast (native) longitudinal</u> <u>relaxation rate</u>, <u>R_10_</u>. Acquisition parameters remained consistent with the dynamic scan except that only six signal averages were acquired, giving a total acquisition time of 46 s per <u>flip angle</u>. To correct <u>R_10_</u> for B_1_ field inhomogeneities and flip angle errors, a pair of images were also acquired from which an actual flip angle map was calculated,^67^ with the same voxel size and coverage as the variable flip angle images. This consisted of a pair of 3D T_1_-FFE images with <u>repetition times</u> of 25 ms and 125 ms, respectively, echo time of 5 ms, a <u>flip angle</u> of 60 degrees, and an acquisition time of 86 s.

A T_1_-weighted image was acquired using an inversion-prepared 3D FFE sequence with turbo factor 204, inversion time 950 ms, repetition time of 6.7 ms, echo time of 3 ms, flip angle 8 degrees, an isotropic resolution of 1 mm^3^, and an acquisition time of 143 s, achieved using compressed sensing factor 4.5. For the assessment of white matter lesions, a T_2_-weighted FLAIR image was acquired with a repetition time of 4800 ms, an inversion time of 1650 ms, echo time of 340 ms, an in-plane resolution of 0.86 mm x 0.86 mm with 150 contiguous axial slices of 1 mm thickness, and an acquisition time of 163 s, achieved using compressed sensing factor 10.

### DCE-MRI image processing & tracer kinetic modelling

A B_1_-corrected pre-contrast (native) R_1_ map (R_10_) map was estimated using the variable flip angle method by fitting the spoiled gradient recalled echo model described in equation 1 to the signal from each voxel of the variable flip angle images using non-linear least-squares minimisation in Madym^68^ v4.23.0.

The dynamic series of post-contrast T_1_-weighted images were aligned using the co-registration function in SPM12 with the first dynamic image as a reference. The first dynamic image was discarded to avoid equilibrium effects. Using an estimated <u>S_0_</u> from the remaining 6 pre-bolus dynamic images, dynamic estimates of <u>R_1_(t)</u> were computed for each voxel of the registered dynamic series of T_1_-weighted images using equation 1. The <u>signal time-course, S_t_(t)</u>, was thus converted to an <u>indicator concentration time-course, C_t_(t)</u>, using the <u>R_10_ map</u> and the <u>longitudinal</u> <u>relaxivity</u> of the contrast agent, which was assumed to be 3.4 s^-1^mM^-1,69^ as described in equation 2. <u>C_t_(t)</u> measurements were also corrected for errors in the applied flip angle using the flip angle map.

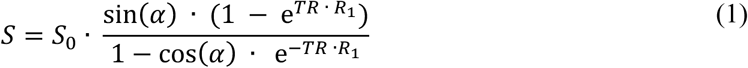

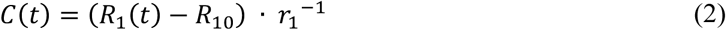

The <u>indicator concentration in plasma, *C*_p_(t)</u>, was calculated using equation 3:

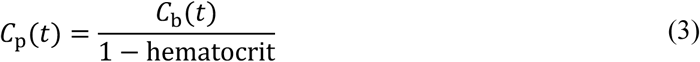

where the <u>indicator concentration in blood, *C*_b_(t)</u>, the vascular input function, was generated using a region <u>drawn manually</u> within the superior sagittal sinus on the <u>last dynamic image</u> in the DCE-MRI series which was eroded to reduce partial volume effects, leaving a region of approximately 10 voxels. The superior sagittal sinus has been recommended by consensus guidelines as it is a large vessel with clear post-contrast enhancement, leading to ease of delineation without partial volume effects, and without inflow effects,^70^ and has been shown to reduce measurement variability compared to an arterial input function.^71^ The mean_signal within this region was extracted for each dynamic to generate *<u>C</u>*<u>_b_(t)</u>. The measured blood <u>hematocrit</u> at the time of the DCE-MRI scan was used for each participant.

For quantification of parameters relating to BBB permeability, a <u>Patlak model</u>^72^ of indicator uptake (equation 4) was fit to the <u>indicator concentration time-course *C*(t)</u> on a voxel-wise basis using constrained <u>non-linear least-squares minimisation</u> in Madym^3^ for 3 parameters: the contrast agent volume transfer constant across the BBB, K^trans^, the <u>fractional blood plasma volume, *v*_p_</u>, and *T*_0_ (where *T*_0_ is the offset time between *<u>C</u>*<u>_t_(t)</u> and *<u>C</u>*<u>_p_(t)</u>). Madym completes this optimisation using Alglib’s boundary and linear equality-inequality constraints algorithm.

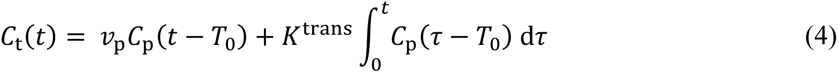

Constraints on the fitted parameters were as follows: between −10 min^-1^ and 10 min^-1^ for K^trans^, between <u>0</u> and <u>(1 – hematocrit)</u> for <u>v_p_</u>, and between <u>-20</u> and <u>20 s</u> for T_0_. Though unphysiological, the negative lower bound for K^trans^ is necessary to avoid positive bias in noisy K^trans^ values, which may be very close to zero in healthy tissue. The negative lower bound on T_0_ allows for the possibility that the regional blood circulation may occur later than the sagittal sinus peak concentration. To reduce errors associated with attempting to fit these models to the rapid concentration changes during the first pass of the contrast agent, the residuals of the first 17 time-points were ignored in the voxel-wise model fitting, corresponding to approximately 70 seconds after the injection of the contrast agent, with each model fit to the remaining 140 time-points.^70,73^ In voxels where the estimated <u>v_p_</u> < 0.1%, the tissue was presumed to be poorly perfused (e.g. inside the ventricles), and these voxels were excluded from further analysis.

#### Post–processing and region of interest analysis

Where possible, the infarcted tissue was defined on imaging as hyperintense signal on a standard clinical diffusion weighted image (DWI) acquired within 14 days of stroke onset. The DWI lesion was delineated manually (ZB) and checked by an experienced stroke neurologist (CS) and neuroradiologist (OMT). DWI collected after 14 days since stroke onset (N = 10) were assessed on a case-by-case basis; where delineation on DWI was considered inappropriate the acute CT and FLAIR were also used to define the infarct region (N = 5). To evaluate changes in the tissue immediately surrounding the acute infarct, the lesion mask was dilated by 1cm using Scipy’s ndimage.binary_dilation to compute a peri-lesion tissue mask. The early DWI/FLAIR was co-registered to the 6-month 3D *T*_1_-weighted image using the co-registration function in SPM12 and the transformation applied to the infarct lesion and peri-lesion masks. Hyperintensities on the 6- month T_2_-weighted FLAIR image, which may consist of small vessel disease white matter lesions or areas of persistent tissue damage from the acute stroke, were defined automatically using the lesion segmentation toolbox (LST)^74^ in SPM12 with the 3D *T*_1_-weighted image as a reference for co-registration, with voxels overlapping with the acute DWI lesion removed from the mask. To avoid ambiguity between the acute and 6-month tissue masks, the acute infarct mask is referred to as the “DWI lesion” region, the dilated acute infarct mask as the “peri-lesion” region, and the 6-month hyperintense FLAIR region as the “white matter hyperintensities” region. Regions of interest corresponding to the normal-appearing gray and white matter were segmented on the 3D *T*_1_-weighted image using SPM’s ‘segment’ function with a threshold of 0.95. The gray and white matter masks were combined to create a “whole brain” mask, and voxels overlapping with the DWI lesion, peri-lesion, or white matter hyperintensities regions removed to generate a mask of the “normal-appearing tissue”, without any lesions.

The <u>first dynamic image</u> of the DCE-MRI acquisition was co-registered to the 3D *T*_1_-weighted image using the co-registration function in SPM12 and the transformation applied to all parametric maps. All co-registered images were visually checked for alignment. The <u>median value</u> of <u>*K*^trans^</u> and *<u>v</u>*<u>_p_</u> was computed for each region in each person.

### Immunostaining and analysis of human brain samples

This study was blinded for all experimental steps including immunostaining, image acquisition, and analysis.

For fluorescent double-staining of endothelial cells (CD31) and mural cells (PDGFRB), human brain sections were deparaffinized in xylene, immersed in decreasing concentrations of ethanol, and rehydrated in water. Antigen retrieval was conducted by subjecting the sections to microwave boiling for 20 minutes in citrate buffer (pH 6). Subsequently, tissue permeabilization is achieved through two 10-minute washes with a buffer containing 1% animal serum and 0.3% Triton X-100 in PBS (PBS-T). The sections were then blocked by incubating them with 10% animal serum in PBS for 1 hour at room temperature. After a brief PBS rinse, autofluorescence was mitigated using an autofluorescence blocker TrueBlack (Biotium) for 30 seconds following the manufacturer’s protocol. After washing with PBS three times, the sections were incubated overnight at 4°C with primary antibodies in blocking buffer. Mouse anti-human PDGFRB (Abcam Ab69506) and rabbit anti-human CD31 (Novus Biologicals NB100-2284) were both diluted at 1:100. Then, sections were rinsed in PBS and exposed to Alexa Fluor 488 and/or 555 conjugated secondary antibodies diluted 1:500 in 1xPBS for 2 hours at room temperature. Finally, after washing 3 times in PBS, sections were mounted using Vectashield mounting medium containing DAPI (Invitrogen). Fluorescence pictures were captured using a Keyence BZ-X700 microscope at 20X magnification.

Three images of each section were taken at random within the sections, then analyzed by FIJI (ImageJ).^75^ The outline of each vessel was manually traced in the CD31 channel to create a binary mask. After denoising the mask using the despeckle extension and removing outliers, the vessel number and area were measured. The CD31 mask was then applied to the PDFGB channel, and the area percentage of overlap between the PDGFB signal and the CD31 mask is measured to quantify mural cell coverage.

Immunostaining of B and T lymphocytes was performed as described.^76^ We used 1:500 CD20 monoclonal mouse anti-human Dako Clone L26 (Agilent t#M0755, RRID:AB_2282030); 1:100 CD3 monoclonal mouse anti-human Dako Clone F7238 (Abcam #ab17143, RRID:AB_302587). Cells were counted manually by a blinded investigator, then normalized to section area to calculate the count per cm^2^.

### Statistical analysis

To compare PDGFB levels between different groups (Stroke vs. Healthy Controls, Better vs. Worse cognitive trajectory) and to adjust for demographic and cardiovascular risk factors, we ran univariate analysis of variance analysis (ANOVA) via General Linear Modeling, where we used log-transformed PDGFB concentrations as the dependent variable.

To test for differences in median *K*^trans^ between stroke survivors and controls in the whole brain and white matter hyperintensities region of interest, respectively, we used Mann-Whitney U tests. To assess differences in *K*^trans^ between the normal-appearing tissue of control brains versus the normal-appearing tissue, stroke lesion, and peri-lesion of stroke survivors, we conducted nonparametric ANOVA using a Kruskall Wallis test with Dunn’s posthoc. To test for paired differences in *K*^trans^ between pathological regions of interest (stroke lesion, peri-lesion, and white matter hyperintensities) and the normal-appearing brain tissue of stroke survivors, we used Wilcoxon signed rank tests.

We conducted an analysis of covariance (ANCOVA) to assess if pericytes coverage is different between infarct groups with and without dementia. As the distribution of the parameters was not normal, we specifically employed Quade’s Nonparametric ANCOVA test of equality of conditional population distributions. We adjusted comparison between groups for age, sex, and presence of atherosclerosis. We investigated if B-cell numbers (CD20 density) mediate the effect of pericyte coverage on cognition category. For this, we followed Baron and Kenny’s 4-step Method for mediation, as well as SAS CAUSALMED procedure, which estimates causal mediation effects from observational data. While conducting 4-step assessment of mediation, we looked into the relationship between pericyte coverage and CD20 as independent variables and dementia as outcome using binary logistic regression. We also checked interaction between independent variables. For non-adjusted comparisons of above variables between groups of interest, we use non-parameteric Mann-Whitney U tests for two groups and Kruskal-Wallis for more than 2 groups. All statistical analyses were 2-sided and significantly different at α=0.05. Statistical software used: Prism 10, SAS 9.4, and IBM SPSS Statistics v30.

## Acknowledgements/Grants

M.S.B. discloses support for this work from the American Heart Association and Allen Frontiers group [AHA 10GRNT4140073] and the Knight Initiative for Brain Resilience. M.S.B, S.A, L.P., and C.J.S. disclose support from the Leducq Foundation [19CVD01] and NINDS NIH [R01NS124927]. V.J.H discloses support from NIH NIA ADRC [P50AG047366 and P30AG066515], B.C.M from NIA NIH [R01AG074339 and R01AG074339], and J.S and A.K from NIH NIA [P30AG072975 and R01AG017917], O.A.J. discloses support for the research of this work from the UKRI EPSRC. K.L. P. discloses support from the Michael J. Fox Foundation. P.C discloses support from the Foundation for Anesthesia Education and Research.

## Author contributions

L.D., J.G., S.M.A, M.L., T.W.C., A.K., J.S., C.J.S., L.M.P., N.A., M.S.B. contributed to conception of the study. L.X., O.A.J., L.Z., N.S.C., E.C.S., E.H., K.B., K.P., E.C.M., A.K., and J.S. contributed to acquisition of the data. L.X., O.A.J., L.D., L.Z., M.M., C.-H.S, Z.B., D.S., O.M.T., P.C., A.A., R.X., J.E.R., H.S.-H.O., L.K.Y., T.J., M.G., P.M.L., K.P., V.W.H., J.G., S.M.A., M.L., E.C.M., T.W.C., C.J.S, L.M.P., N.A., and M.S.B. contributed to analysis of the data. L.X., O.A.J., L.D., L.Z., M.M., C.-H.S., and M.S.B contributed to interpretation of the data. L.X., O.A.J., L.D., K.Z., L.Z., M.M., P.M.L., S.M.A., E.C.M., T.W.C., A.K., J.S., C.J.S, L.M.P., N.A., and M.S.B. drafted the manuscript. All authors reviewed and approved the submitted version of the manuscript.

## Additional information

Correspondence and requests for materials should be addressed to Dr. Marion Buckwalter at.

## Data Availability

The datasets generated during and/or analysed during the described studies are available from the corresponding author on reasonable request.

## Code Availability

Code is available at GitHub (https://github.com/leixuecynthia/StrokeCog-analysis) and archived at Zenodo (DOI: 10.5281/zenodo.18677368).

