## Supplemental Figures and Tables for "Blood-brain barrier dysfunction predicts cognitive trajectory after ischemic stroke"

**Additional information**

**Data Availability**

The datasets generated during and/or analysed during the described studies are available from the corresponding author on reasonable request.

**Code Availability**

Code is available at GitHub (https://github.com/leixuecynthia/StrokeCog-analysis) and archived at Zenodo (DOI: 10.5281/zenodo.18677368).

### Extended Data Figures

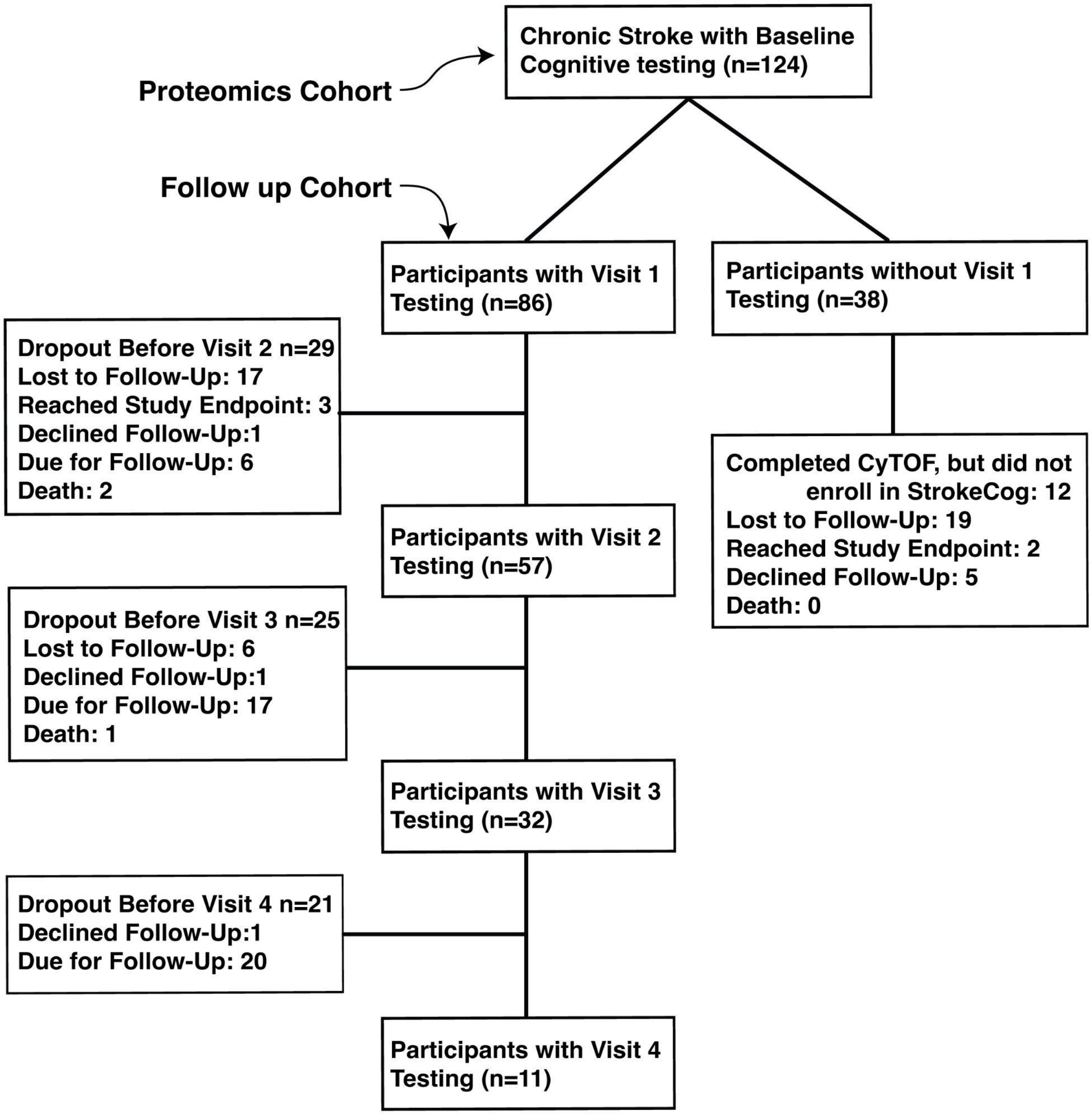


**Extended Data Figure 1. Flowchart of participants in StrokeCog.** CyTOF was an earlier study of immunophenotypes after stroke[42] and participants were offered the chance to continue follow-up as part of the StrokeCog study. Average follow-up in the cohort was 26 months at the time of analysis. Note that many who were lost to follow up for Year 1 and 2 visits were lost during the COVID-19 pandemic. Participants that were "due for follow-up" at the time of data analysis were still in the study but not yet in window for their next cognitive follow-up. Reasons for reaching study endpoint included MoCA<11 or inability to perform cognitive testing due to cognitive issues.

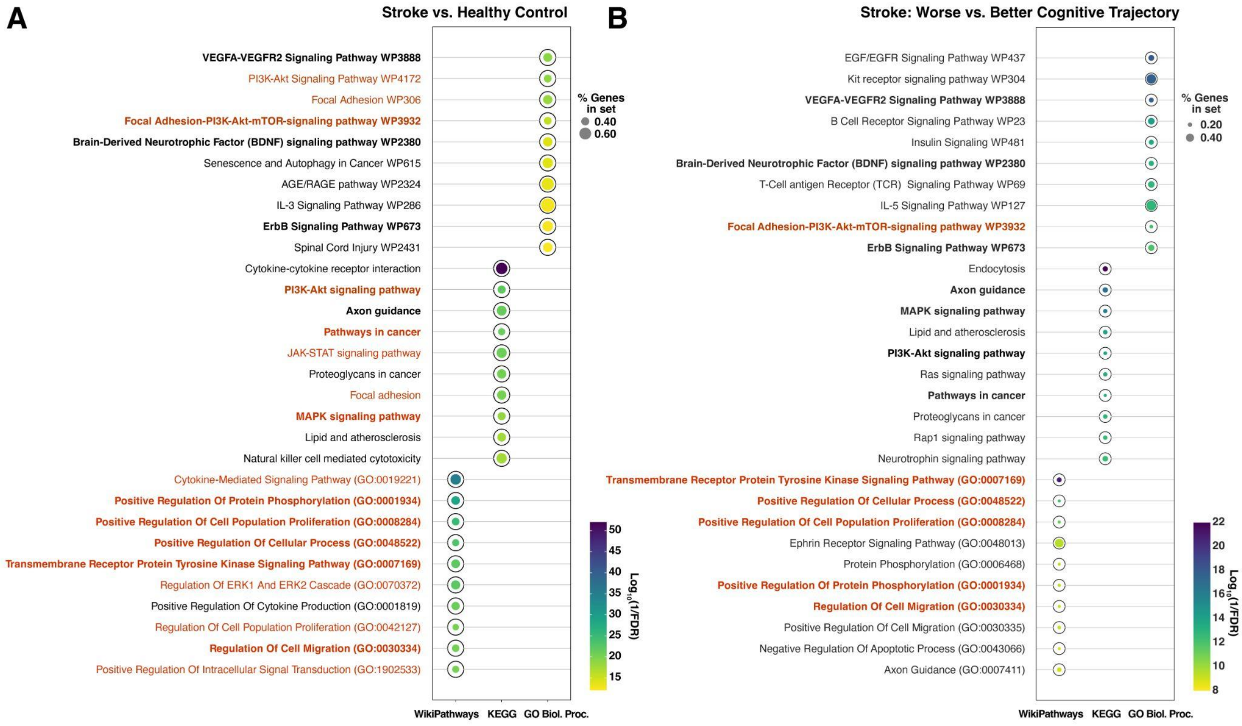


**Extended Data Figure 2. Over-representation analysis of plasma proteomics.** Bolded pathways appear in both comparisons and red lettering highlights pathways that are driven by changes in PDGFB and its related genes. Top ten by FDR, from top to bottom, Wiki Pathways, KEGG, and GO Pathways.

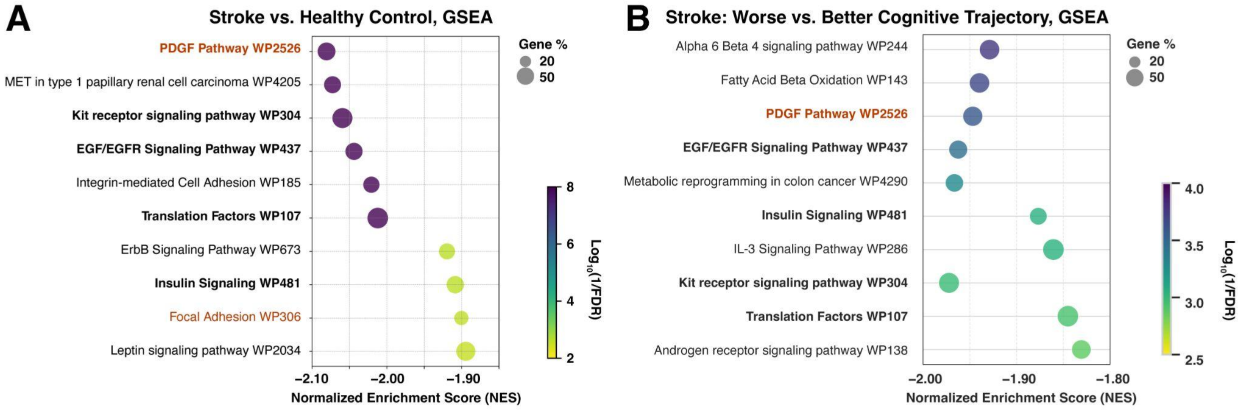


**Extended Data Figure 3. Gene set enrichment analysis of plasma proteomics.** Bolded pathways appear in both comparisons and red lettering highlights pathways that are driven by changes in PDGFB and its related genes.

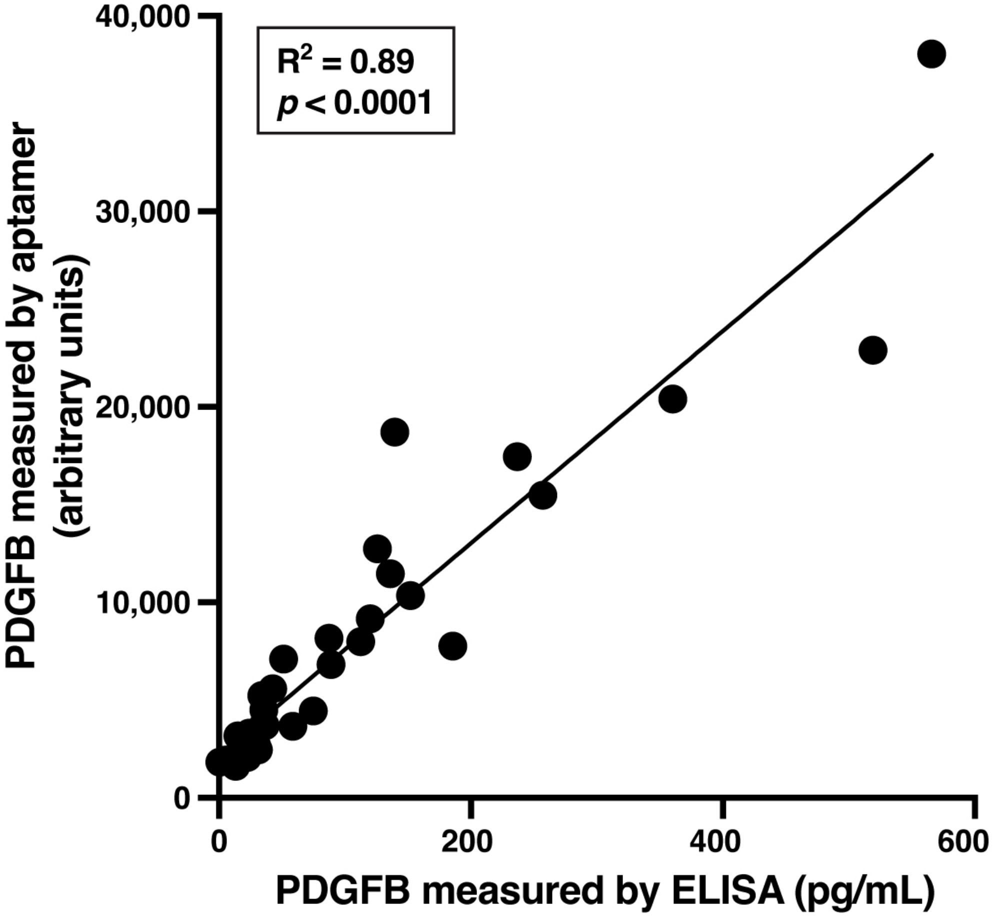


**Extended Data Figure 4. PDGF-BB measured by aptamer correlated well with ELISA measures.** Linear regression.

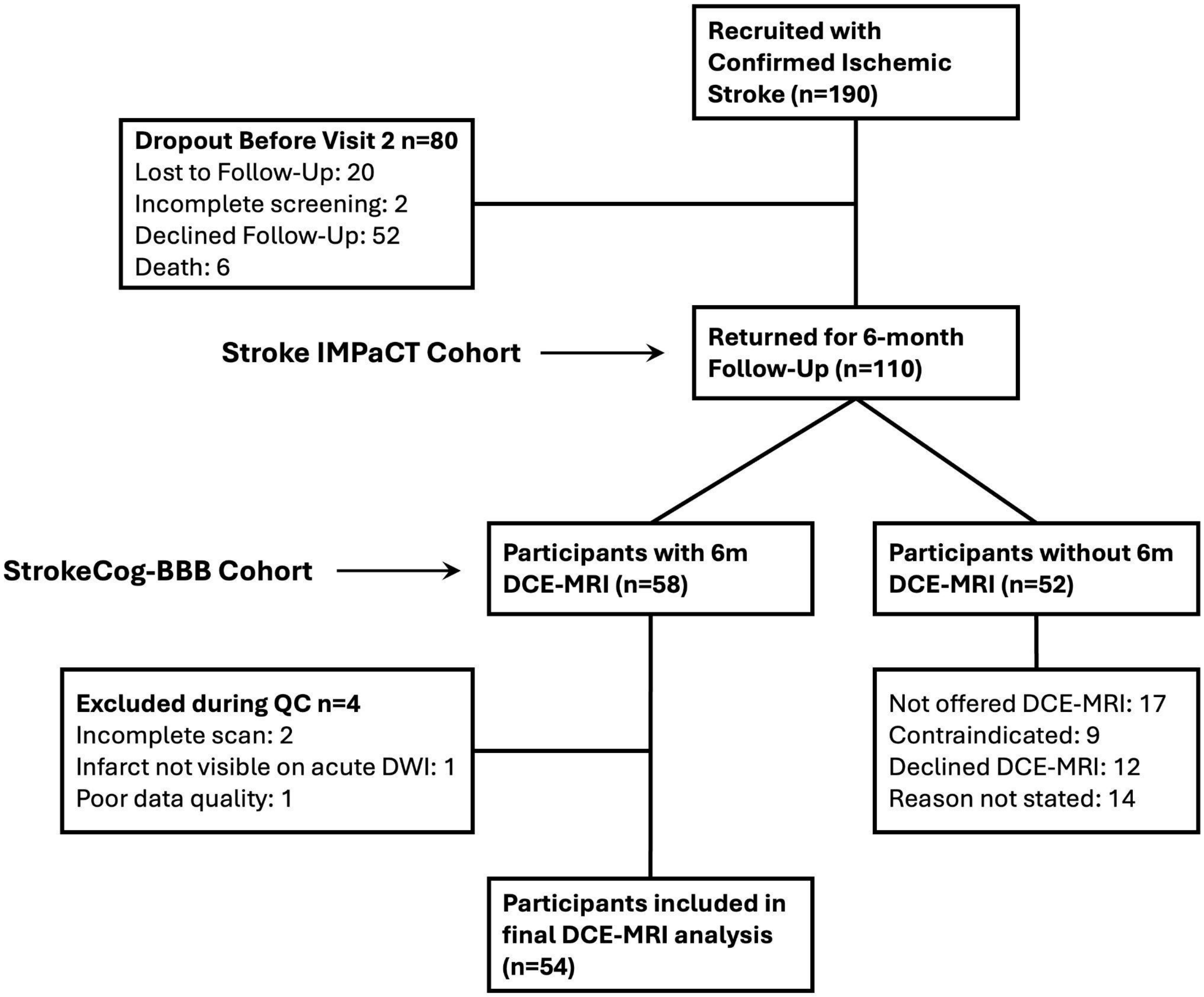


**Extended Data Figure 5. Flowchart of participants in StrokeCog-BBB.** Participants in the Stroke IMPaCT study were offered DCE-MRI at the 6 month time-point. Seventeen participants were recruited before the StrokeCog-BBB study began and were not offered DCE-MRI, 9 participants were contraindicated to DCE-MRI due to low or absent eGFR, non-MR compatible devices, etc.

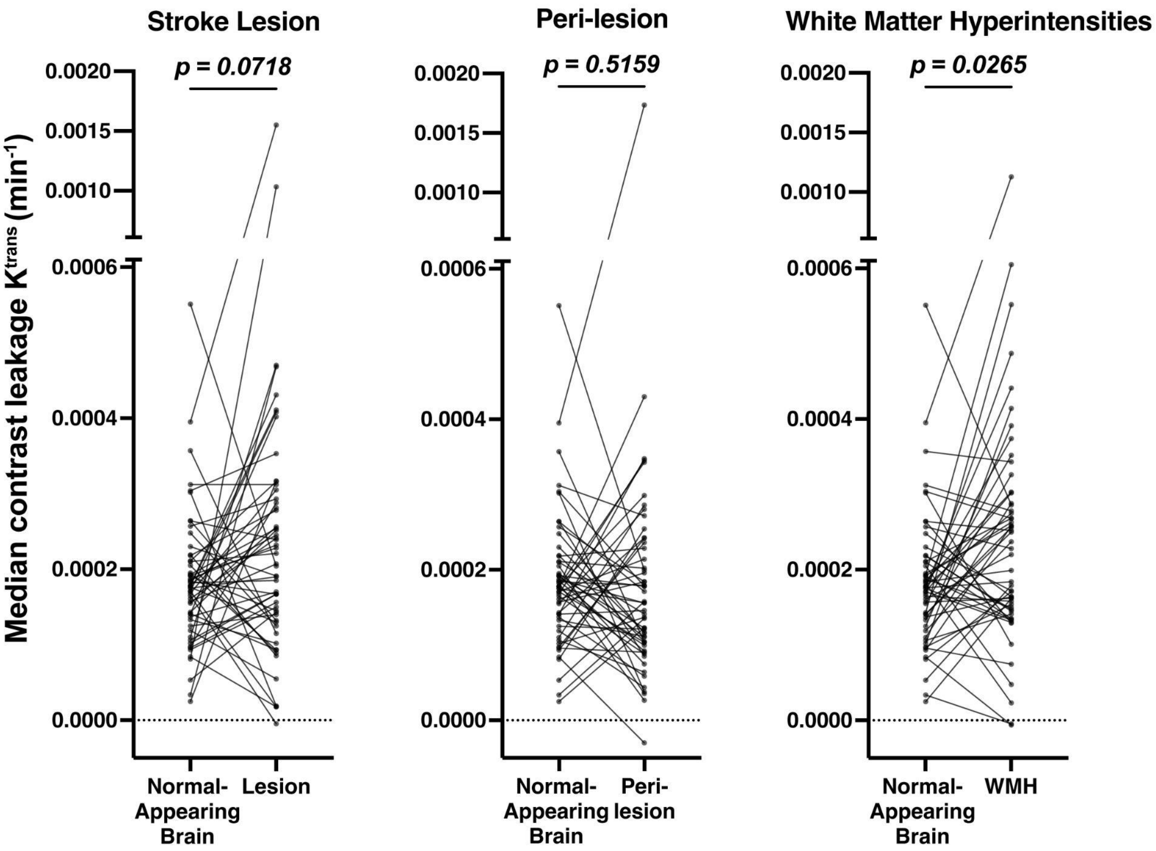


**Extended Data Figure 6. Comparison of $K^{trans}$ between brain regions in chronic stroke.** Paired comparison of median $K^{trans}$ estimates in the normal-appearing tissue vs. old stroke lesion, peri-lesion, and white matter hyperintensities of stroke participants. *p*-values, Wilcoxon matched-pairs signed rank test.

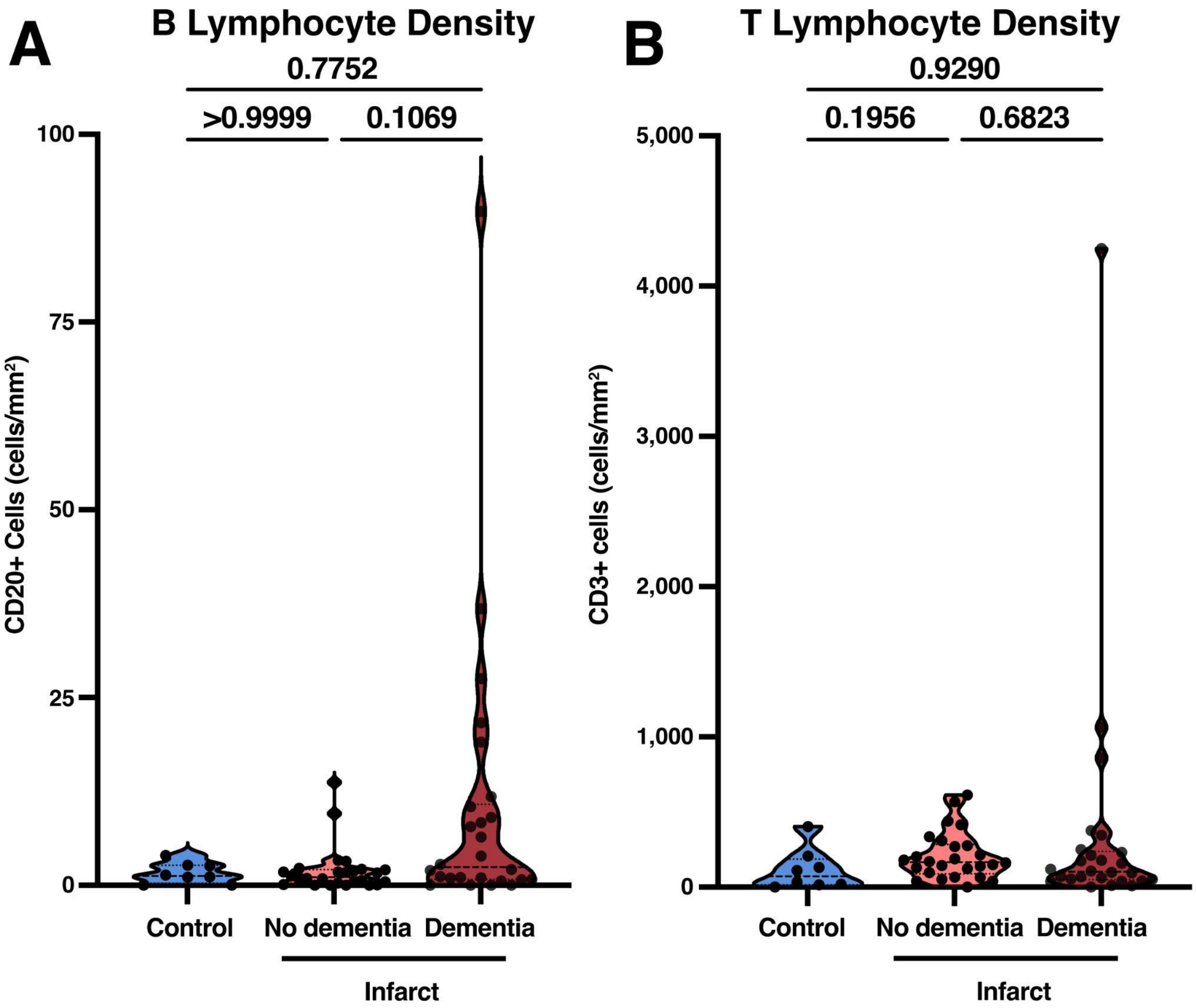


**Extended Data Figure 7. Lymphocyte densities in infarcts from the autopsy cohort. A)** Quantification of CD20-expressing B lymphocyte density in those who died with infarcts and dementia, infarcts and no dementia, and controls. **B)** Quantification of CD3-expressing T lymphocyte density in those who died with infarcts and dementia, infarcts and no dementia, and controls. n=8 for controls and n=26 for each infarct group. *p*-values, Kruskal-Wallis non-parametric ANOVA with Tukey's post-hoc.

**Extended Data Tables**

| Aptamer Name | Gene | Protein Name | EN model Coefficient |
|---|---|---|---|
| C1R.3285.23.2 | C1R | Complement C1r subcomponent | -0.820 |
| C4A.C4B.18821.9.3 | C4A.C4B | C4a anaphylatoxin | -0.499 |
| SCIN.12684.5.3 | SCIN | Adseverin | 0.479 |
| CPQ.9394.19.3 | CPQ | Carboxypeptidase Q | -0.469 |
| SHH.2743.5.2 | SHH | Sonic hedgehog protein | 0.431 |
| C4A.C4B.2182.54.1 | C4A.C4B | Complement C4b | -0.385 |
| IGFBP1.2771.35.2 | IGFBP1 | Insulin-like growth factor-binding protein 1 | -0.369 |
| IHH.19606.28.3 | IHH | Indian hedgehog protein | 0.338 |
| NIF3L1.23371.5.3 | NIF3L1 | NIF3-like protein 1 | -0.333 |
| GLIPR2.15522.2.3 | GLIPR2 | Golgi-associated plant pathogenesis-related protein 1 | -0.333 |
| HNF4A.10041.3.3 | HNF4A | Hepatocyte nuclear factor 4-alpha | 0.284 |
| ERP29.4983.6.1 | ERP29 | Endoplasmic reticulum resident protein 29 | 0.284 |
| CFD.13678.169.3 | CFD | Complement factor D | 0.280 |
| KNG1.7784.1.3 | KNG1 | Kininogen-1 | -0.277 |
| ENPP2.16892.23.3 | ENPP2 | Ectonucleotide pyrophosphatase/phosphodiesterase family member 2 | 0.252 |
| GCG.4891.50.1 | GCG | Glucagon | 0.235 |
| NXPH1.4562.1.2 | NXPH1 | Neurexophilin-1 | -0.230 |
| IGFBP1.13741.36.3 | IGFBP1 | Insulin-like growth factor-binding protein 1 | -0.225 |
| ARPC2.23376.56.3 | ARPC2 | Actin-related protein 2/3 complex subunit 2 | -0.217 |
| TOM1L1.13652.2.3 | TOM1L1 | TOM1-like protein 1 | -0.212 |

**Extended Data Table 1. Top 20 Features in the Elastic Net Model.** List of the 20 most important features based on the absolute value of their coefficients from the Elastic Net regression model. Feature importance reflects the relative contribution of each feature to the model's predictive performance. Positive coefficients indicate features associated with an increase in the target outcome, while negative coefficients indicate features associated with a decrease.

| Gene Name | Aptamer Name | Protein Names (Name if used in Fig 2) | Member of an annotated protein community | $Log_2FC$ | Corre-lation | F-Statistic |
|---|---|---|---|---|---|---|
| **GIP** | GIP.16292.288.3 | Gastric inhibitory polypeptide, aa 1-93 (GIP) | Yes | 0.82 | 0.99 | 282.28 |
| | GIP.5755.29.3 | Gastric inhibitory polypeptide, aa 22-153 | N/A | 0.09 | 0.87 | 32.50 |
| **MMP8** | MMP8.2954.56.2 | Neutrophil collagenase, aa 21-467 | N/A | 0.03 | 0.88 | 9.23 |
| | MMP8.9172.69.3 | Neutrophil collagenase, aa 21-467 (MMP8) | Yes | 0.93 | 0.97 | 7.96 |
| **MMP7** | MMP7.2789.26.2 | Matrilysin, aa 18-267 (MMP7) | Yes | 1.09 | 0.89 | 16.85 |
| | MMP7.8475.15.3 | Matrilysin, aa 95-267 | N/A | 0.04 | 0.55 | 3.01 |
| **PCK2** | PCK2.20079.6.3 | Phosphoenolpyruvate carboxykinase [GTP], mitochondrial, aa 33-198 | N/A | -0.06 | 0.38 | 2.75 |
| | PCK2.22405.61.3 | Phosphoenolpyruvate carboxykinase [GTP], mitochondrial, aa 33-640 (PCK2) | Yes | 1.43 | 0.97 | 26.71 |
| **KNG1** | KNG1.15343.337.3 | Kininogen, HMW, Two Chain (KNG1 (a)) | Yes | 1.23 | 0.69 | 4.83 |
| | KNG1.19631.13.3 | Kininostatin (KNG1 (b)) | Yes | 2.04 | 0.90 | 16.13 |
| | KNG1.4918.21.1 | Kininogen-1 | N/A | 0.00 | 0.17 | 1.54 |
| | KNG1.7784.1.3 | Kininogen-1 | N/A | -0.31 | 0.64 | 49.90 |
| **FOLH1** | FOLH1.3218.8.2 | Glutamate carboxypeptidase 2, aa 44-750 | N/A | 0.00 | 0.67 | 7.00 |
| | FOLH1.5478.50.2 | Glutamate carboxypeptidase 2, aa 44-750 (FOLH1) | Yes | 1.51 | 0.98 | 413.94 |
| **ALPI** | ALPI.10463.23.3 | Intestinal-type alkaline phosphatase, aa 20-528 | N/A | 0.04 | 0.76 | 6.45 |
| | ALPI.17441.4.3 | Intestinal-type alkaline phosphatase, aa 1-503 (ALP1) | Yes | 0.88 | 0.87 | 48.17 |
| **C4A** | C4A.C4B.18821.9.3 | C4a anaphylatoxin (C4A.C4B (a)) | Yes | -2.32 | 0.95 | 36.45 |
| | C4A.C4B.2182.54.1 | Complement C4b (C4A.C4B (b)) | Yes | -0.60 | 0.88 | 9.27 |
| | C4A.C4B.4481.34.2 | Complement C4 | N/A | 0.12 | 0.73 | 5.06 |
| **CCL5** | CCL5.2523.31.3 | C-C motif chemokine 5, aa 24-91 (CCL5 (a)) | Yes | -0.80 | 0.94 | 51.66 |
| | CCL5.5480.49.3 | C-C motif chemokine 5 (CCL5 (b)) | Yes | -1.20 | 0.93 | 91.04 |
| **PDGFD** | PDGFD.17140.57.3 | Platelet-derived growth factor D, aa 250-370 (PDGFD) | Yes | -1.00 | 0.98 | 112.58 |
| | PDGFD.9341.1.3 | Platelet-derived growth factor D, aa 19-364 | N/A | -0.18 | 0.99 | 175.96 |

| Gene Name | Aptamer Name | Protein Names (Name if used in Fig 2) | Member of an annotated protein community | $Log_2FC$ | Corre-lation | F-Statistic |
|---|---|---|---|---|---|---|
| **SERPINE2** | SERPINE2.19154.41.3 | Glia-derived nexin, aa 1-397 (SERPINE2) | Yes | -0.84 | 0.98 | 72.73 |
| | SERPINE2.3217.74.2 | Glia-derived nexin, aa 1-397 | N/A | -0.20 | 0.94 | 2.05 |
| **DSG3** | DSG3.11310.8.3 | Desmoglein-3, aa 24-615 | N/A | 0.08 | 0.77 | 3.73 |
| | DSG3.16317.20.3 | Desmoglein-3, aa 50-615 (DSG3) | Yes | -0.98 | 0.99 | 219.78 |
| **BDNF** | BDNF.14047.78.3 | Brain-derived neurotrophic factor (BDNF (a)) | Yes | -1.11 | 0.98 | 31.91 |
| | BDNF.2421.7.3 | Brain-derived neurotrophic factor (BDNF (b)) | Yes | -0.48 | 0.96 | 4.82 |
| **PPBP** | PPBP.16765.52.3 | Platelet basic protein (PPBP (a)) | Yes | -0.66 | 0.97 | 118.87 |
| | PPBP.2790.54.2 | Neutrophil-activating peptide 2 (PPBP (c)) | Yes | -1.21 | 0.97 | 24.24 |
| | PPBP.17165.1.3 | Beta-thromboglobulin (PPBP (b)) | Yes | -1.05 | 0.98 | 26.21 |
| | PPBP.4544.4.3 | Connective tissue-activating peptide III (PPBP (d)) | Yes | -1.22 | 0.97 | 33.20 |

**Extended data table 2. Aptamers in significant protein communities that have alternate targets.** This table presents detailed information about the SomaScan aptamers included in the protein communities of interest where there is more than one targeting aptamer, and their corresponding proteins selected in the analysis, along with key statistical metrics. **$Log_2FC$** represents the $\log_2$ (fold change) between Stroke and Healthy Control groups, so a negative number means there is a lower protein concentration in the Stroke group in our analysis. The next three values are from SomaLogic's quality control measures. **Correlation** is the average run-to-run correlation coefficient for this SOMAmer reagent in repeated tests of human plasma from healthy individuals. **F-statistic** is the ratio of signal variance from a set of healthy individuals divided by signal variance from technical replicates. F-statistics above the critical value (2.43) indicate measured biological differences during testing of the aptamers in human samples by the company, so higher values mean higher ability to detect biological differences in our samples.

| | ADRC Healthy Controls (N = 187) | Stroke (N = 124) |
|---|---|---|
| **Age (years)** | 72 [68, 77] | 65 [55, 74]*** |
| **Sex (% Female)** | 61% | 45%** |
| **Medications** | | |
| Hypertension | 34% | 52%** |
| Hyperlipidemia | 43% | 66%*** |
| Diabetes | 4.3% | 13%** |
| **PDGFB by aptamer (arbitrary units)** | | |
| All | 19,215 [10,270, 33,787] | 6,421 [2,563, 16,462]*** |
| No Hypertension (n = 123, 59) | 18,369 [9,827, 27,802] | 6,757 [2,379, 16,902]*** |
| Hypertension (n = 64, 65) | 22,167 [11,186, 39,767] | 6,079 [2,658, 16,353]*** |
| No Hyperlipidemia (n = 106, 42) | 18,695 [9,498, 33537] | 5,896 [2,344, 9,818]*** |
| Hyperlipidemia (n = 81, 82) | 19,599 [11,225, 35,362] | 7,124 [2,793, 19,619]*** |
| No Diabetes (n = 180, 108) | 19,003 [10,265, 33,665] | 6,518 [2,628, 16,462]*** |
| Diabetes (n = 7, 16) | 26,200 [18,364, 38,093] | 4,069 [1,842, 15,769]** |
| No Cardiovascular risk factors (n = 83, 28) | 16,764 [9,008, 27,753] | 6,674 [2,275, 18,837]** |
| 1 Cardiovascular risk factor (n = 60, 39) | 21,999 [11,413, 36,534] | 6,079 [2,717, 11,462]*** |
| 2 Cardiovascular risk factors (n = 39, 47) | 16,586 [11,147, 36,937] | 5,945 [1,947, 17,876]*** |
| 3 Cardiovascular risk factors (n = 5, 10) | 33,462 [16,467, 114,871] | 8,145 [3,272, 25,950] |

**Extended Data Table 3. Effect of cardiovascular risk on PDGFB aptamer levels in ADRC Healthy Control participants and Stroke participants.** These are unmatched datasets of all healthy controls with cardiovascular risk available, all from Stanford's Alzheimer's Disease Research Center, and all the Stanford Stroke participants. Demographics are listed including three cardiovascular risk factors, defined as those on medication for hypertension, hyperlipidemia and diabetes. For Age and PDGFB, *p*-values were calculated using the Mann-Whitney U test. For Sex and Medications, *p*-values were calculated using the Chi-squared test. *, Comparison between Controls and Stroke participants. **$p<0.01$; ****$p<0.001$. For cardiovascular risks, there are no differences in PDGFB between any category within control or stroke participants. Between ADRC Controls and Stroke, *p*-values were calculated using the Mann-Whitney U test.

| Variable | Beta (95% confidence interval) | Partial Eta Squared | *p*-value |
|---|---|---|---|
| **Stroke** | -1.154 (-1.403 - -0.906) | 0.215 | <0.001 |
| **Age** | -0.00007 (-0.011 - 0.011) | 0.0000006 | 0.990 |
| **Sex (% Female)** | 0.049 (-0.173 - 0.271) | 0.001 | 0.664 |
| **Hypertension** | 0.048 (-0.194 - 0.290) | 0.001 | 0.696 |
| **Hyperlipidemia** | -0.067 (-0.486 - 0.352) | 0.0003 | 0.754 |
| **Diabetes** | 0.147 (-0.090 - 0.384) | 0.005 | 0.223 |

**Extended Data Table 4.** Univariate ANOVA general linear model to test whether the natural log of PDGFB by aptamer measure (arbitrary units) is dependent on the listed variables. Partial Eta squared demonstrates that 22% of the variance is due to stroke, which is the only significant variable. B, beta, Partial Eta squared, percent variance due to that variable.

| Demographics | All Controls (N = 15) | All Stroke (N = 54) | Stroke Low Leakage (N = 27) | Stroke High Leakage (N = 27) |
|---|---|---|---|---|
| **Age (years)** | 69 [60, 72] | 65 [57, 71] | 68 [60, 71] | 64 [57, 69] |
| **Sex (% Male)** | 73% | 82% | 85% | 78% |
| **Race** | | | | |
| White | 93% | 96% | 93% | 100% |
| Asian | 7% | 4% | 7% | 0% |
| **Medications** | | | | |
| Hypertension | 73% | 59% | 44% | 74%* |
| Hyperlipidemia | 80% | 89% | 82% | 96% |
| Diabetes | 47% | 19%† | 19% | 19% |
| **Time Since Stroke (days)** | N/A | 206 [195, 225] | 202 [194, 224] | 207 [197, 228] |
| **Stroke Size (mL)** | N/A | 2.1 [0.75, 6.30] | 2.2 [0.75, 10.47] | 1.6 [0.82, 5.46] |
| **Stroke Severity (NIHSS)** | N/A | 3.0 [2.0, 6.0] | 4.0 [2.0, 5.5] | 3.0 [2.0, 7.5] |
| **Cognitive z-score** | | | | |
| Visuospatial | 1.00 [0.67, 1.67] | 1.00 [0.00, 1.33] | 1.00 [0.34, 1.21] | 1.00 [0.08, 1.33] |
| Memory | -0.45 [-1.15, 0.48] | -0.98 [-1.75, -0.05] | -1.05 [-1.85, -0.45] | -0.80 [-1.63, 0.20] |
| Processing speed | 0.27 [-0.33, 0.56] | -0.42 [-0.92, 0.23]† | -0.34 [-0.85, -0.15] | -0.50 [-1.18, 0.21] |
| Working memory | 0.33 [-0.17, 0.84] | 0.00 [-0.33, 1.00] | 0.17 [-0.33, 0.67] | 0.00 [-0.50, 1.00] |
| Language | 0.65 [0.42, 1.52] | 0.29 [-0.35, 0.73]† | 0.20 [-0.41, 0.41] | 0.57 [-0.13, 0.80] |
| **$K^{trans}$ ($10^{-3}$ $min^{-1}$)** | | | | |
| Whole brain | 0.103 [0.037, 0.125] | 0.179 [0.133, 0.210]† | 0.133 [0.096, 0.159] | 0.210 [0.185, 0.267]* |
| Lesion | N/A | 0.206 [0.132, 0.296] | 0.156 [0.093, 0.227] | 0.252 [0.179, 0.314]* |
| Peri-lesion | N/A | 0.151 [0.103, 0.255] | 0.178 [0.140, 0.249] | 0.290 [0.223, 0.372]* |
| White matter hyperintensities | 0.098 [0.077, 0.216] | 0.224 [0.147, 0.292]† | 0.158 [0.130, 0.182] | 0.278 [0.231, 0.363]* |
| Normal-appearing tissue | 0.103 [0.033, 0.133] | 0.177 [0.133, 0.211]† | 0.133 [0.097, 0.159] | 0.211 [0.188, 0.264]* |
| **$v_p$ (%)** | | | | |
| Whole brain | 0.605 [0.413, 0.885] | 0.634 [0.482, 0.723] | 0.621 [0.479, 0.728] | 0.659 [0.485, 0.721] |
| Lesion | N/A | 0.543 [0.407, 0.726] | 0.532 [0.357, 0.753] | 0.562 [0.431, 0.661] |
| Peri-lesion | N/A | 0.617 [0.492, 0.727] | 0.606 [0.477, 0.726] | 0.626 [0.508, 0.742] |
| White matter hyperintensities | 0.6960 [0.469, 1.00] | 0.598 [0.413, 0.819] | 0.602 [0.414, 0.758] | 0.587 [0.411, 0.903] |
| Normal-appearing tissue | 0.605 [0.414, 0.884] | 0.651 [0.484, 0.731] | 0.621 [0.482, 9.735] | 0.660 [0.484, 0.730] |
| **PDGF-BB (pg/mL)** | 522 [427, 628] | 387 [213, 516] | 303 [209, 495] | 437 [267, 519] |
| **Compared to:** | | **All Controls** | | **Stroke Low Leakage** |

**Extended Data Table 5. Demographic and Clinical Characteristics of the DCE-MRI Cohort.** "All Stroke" indicates all stroke participants who had a DCE-MRI scan, Stroke "Low Leakage" and "High Leakage" indicate those in the cohort whose whole brain $K^{trans}$ was lower or higher than the group median. The "All Stroke" cohort has 4 missing values for working memory z-score and 6 missing values for PDGFBB. The Stroke "High Leakage" group has 3 missing values for working memory z-score and 4 missing values for PDGFBB. The Stroke "Low Leakage" group has 1 missing value for working memory z-score and 2 missing values for PDGFBB. The "All Controls" cohort has 8 missing values for PDGFBB. Stroke Severity was measured by National Institutes of Health Stroke Scale, NIHSS. For Age, Stroke Size, Time Since Stroke, NIHSS, Cognitive z-scores, $K^{trans}$, $v_p$, and PDGFBB, $p$-values were calculated using the Mann-Whitney U test. For Sex and Medications, *p*-values were calculated using the Chi-squared test. [†], different between stroke and controls; *, different between low and high leakage groups.

| | Controls (n = 8) | Infarcts and no Dementia (n = 26) | Infarcts and Dementia (n = 26) |
|---|---|---|---|
| **Age (years)** | 87 [84, 94] | 88 [84, 93] | 91 [81, 96] |
| **Sex (% Male)** | 63% | 46% | 23% |
| **Race** | | | |
| White | 100% | 92% | 100% |
| Black | 0% | 4% | 0% |
| Unknown | 0% | 4% | 0% |
| **Post-mortem interval (hours)** | 4.7 [1.6, 7.5] | 7.8 [5.3, 12.8] | 6.1 [4.4, 9.2] |
| **Comorbidities** | | | |
| Hypertension | 50% | 73% | 81% |
| Diabetes | 25% | 31% | 27% |
| BMI | 28 [26, 31] | 30 [26, 33] | 28 [25, 32] |
| **Premorbid Clinical Diagnosis (cogdx)** | | | |
| No cognitive impairment | 62.5% | 46.2% | 0% |
| MCI and no other cause of CI | 25.0% | 50.0% | 0% |
| MCI and another cause of CI | 12.5% | 3.8% | 0% |
| AD and no other cause of CI | 0% | 0% | 42.3% |
| AD and another cause of CI | 0% | 0% | 42.3% |
| Other dementia | 0% | 0% | 15.4% |
| **Brain Pathology** | | | |
| Global AD pathology | 0.08 [0.02, 0.17] # | 0.13 [0.09, 0.50] # | 0.14 [0.08, 0.24] # |
| Amyloid density | 0.05 [0.00, 0.27] | 0.36 [0.00, 0.94] # | 0.05 [0.00, 0.77] # |
| Braak stage | 2.5 [1.0, 3.0] # | 3.0 [2.0, 3.0] # | 3.0 [2.0, 4.0] # |
| CERAD score | 4.0 [3.3, 4.0] # | 4.0 [2.0, 4.0] # | 4.0 [3.0, 4.0] # |
| Arteriosclerosis | 1.0 [0.25, 2.8] # | 1.0 [1.0, 2.0] # | 2.0 [1.0, 2.0] # |
| Atherosclerosis | 1.0 [0.25, 1.8] # | 1.0 [1.0, 2.0] # | 2.0 [2.0, 3.0] †, *, # |
| CAA | 0.5 [0.00, 1.0] # | 1.0 [0.0, 1.0] # | 1.0 [0.0, 2.0] # |
| Number of subcortical lacunar infarcts | N/A | 1.0 [0.0, 2.0] † | 1.0 [0.0, 1.3] † |
| Gross infarcts (yes/no) | N/A | 92.3% | 88.5% |
| Microinfarcts (yes/no) | N/A | 34.6%% | 42.3% |
| **Residual Cognitive Trajectory** (corrected for known brain pathologies) | | | |
| Global | -0.0066 [-0.026, 0.0079] | 0.025 [0.00015, 0.05]# | -0.038 [-0.087,0.00016]*,# |
| Episodic Memory | -0.018 [-0.051, -0.014] | 0.012 [-0.00081, 0.053]# | -0.036 [-0.76, 0.023]*,# |
| Perceptual Speed | 0.017 [-0.010, 0.030] | 0.014 [-0.021, 0.050] | -0.020 [-0.069, -0.0056]*,# |
| Visuospatial/perceptual organization | 0.0015 [-0.0062, 0,0081] | 0.0077 [-0.025, 0.030] | -0.031[-0.065, -0.0080]*,†,# |
| Semantic Memory | -0.0017 [-0.029, 0.021] | 0.017 [-0.019, 0.045] | -0.049 [-0.074, 0.032]*,# |
| Working Memory | -0.0051 [-0.022, 0.016] | 0.011 [-0.0032, 0,033] | -0.028 [-0.058, -0.000005]*,# |

**Extended Data Table 6. Clinical and pathological characteristics of the autopsy cohort.** Clinical and pathological characteristics of the autopsy cohort. Demographic, neuropathological and cognitive data is listed for each group. Premorbid cognitive diagnosis (cogdx) was made by neurologists blinded to pathology using all available clinical data. The overall distribution of cogdx is different between all 3 groups, $p<0.001$; different between infarct with and without dementia, $p<0.001$; with dementia group is different from controls, $p<0.001$, but the without dementia group is not different from controls. MCI, mild cognitive impairment defined as one impaired cognitive domain; CI, cognitive impairment; AD, Alzheimer's Disease. Kruskal Wallis was used for continuous and ordered variables such as pathological scores (e.g. arteriosclerosis is a 4-point scale, 0-3) and Chi squared test for categorical variables, with a one-Sample Wilcoxian signed rank test to ask if values were different from zero. $^{\dagger}$, different from controls; *, different between infarct with and without dementia; $^{\#}$, different from zero.
